# High GC Content Causes Orphan Proteins to be Intrinsically Disordered

**DOI:** 10.1101/103739

**Authors:** Walter Basile, Oxana Sachenkova, Sara Light, Arne Elofsson

## Abstract

*De novo* creation of protein coding genes involves the formation of short ORFs from noncoding regions; some of these ORFs might then become fixed in the populationThese orphan proteins need to, at the bare minimum, not cause serious harm to the organism, meaning that they should for instance not aggregate. Therefore, although the creation of short ORFs could be truly random, the fixation should be subjected to some selective pressure. The selective forces acting on orphan proteins have been elusive, and contradictory results have been reported. In *Drosophila* young proteins are more disordered than ancient ones, while the opposite trend is present in yeast. To the best of our knowledge no valid explanation for this difference has been proposed.

To solve this riddle we studied structural properties and age of proteins in 187 eukaryotic organisms. We find that, with the exception of length, there are only small differences in the properties between proteins of different ages. However, when we take the GC content into account we noted that it could explain the opposite trends observed for orphans in yeast (low GC) and *Drosophila* (high GC). GC content is correlated with codons coding for disorder promoting amino acids. This leads us to propose that intrinsic disorder is not a strong determining factor for fixation of orphan proteins. Instead these proteins largely resemble random proteins given a particular GC level. During evolution the properties of a protein change faster than the GC level causing the relationship between disorder and GC to gradually weaken.

**Author Summary:** We show that the GC content of a genome is of great importance for the properties of an orphan protein. GC content affects the frequency of the codons and this affects the probability for each amino acid to be included in a *de novo* created protein. The codons encoding for Ala, Pro and Gly contain 80% GC, while codons for Lys, Phe, Asn, Tyr and Ile contain 20% or less. The three high GC amino acids are all disorder promoting, while Phe, Tyr and Ile are order promoting. Therefore, random protein sequences at a high GC will be more disordered than the ones created at a low GC. The structural properties of the youngest proteins match to a large degree the properties of random proteins when the GC content is taken into account. In contrast, structural properties of ancient proteins only show a weak correlation with GC content. This suggests that even after fixation in the population, proteins largely resemble random proteins given a certain GC content. Thereafter, during evolution the correlation between structural properties and GC weakens.

## Introduction

Proteins without any detectable homology are often referred to as orphans. The presence of orphans can be attributed to several causes; rapid sequence divergence beyond the point of homology recognition [1, 2], lateral transfer of genetic material [3], and *de novo* gene creation [4]. The latter is of particular interest, as it is a source of completely novel coding material. Studies of the properties of these proteins might provide unique insights into the fundamental processes in the formation of all proteins, since, in the strict sense, all proteins were once created by a *de novo* mechanism.

Before the genomic era, the scientific consensus held that *de novo* creation of new genes was rare - instead it was believed that the vast majority of all genes were generated in an ancient “big bang”. However, when the first complete genomic sequences were initially published, this hypothesis was not supported [5]. In fact, to this day, when analyzing complete genomes from closely related species, a surprisingly high number of orphan proteins is still found [6–8]. It has later been shown that some of these proteins are not *de novo* created but rather assigned as orphans as a result of limited phylogenetic coverage in earlier studies [9].

Today supported by the vast amount of complete genome sequences available and improved search methods [10], many of the initially identified orphans have been shown to have distant homologs in other genomes. Still, at least in yeast, a large set of genes appears to have been created through recent *de novo* formation [11, 12]. These studies indicate that in yeast there is a large set of proto-genes: ORFs that remain on the verge of becoming fixed as protein-coding genes in the population [11]. This provides a possible model of how novel proteins can be generated from noncoding genetic material. In other species than yeast the genomic coverage is more limited and therefore studies have been less detailed.

The availability of many, complete, evenly spaced genomes allows classifying proteins at different evolutionary age [7, 9, 11], using methods such as ProteinHistorian [13]. Here, a protein can be unique to a specific species, or even to a strain; alternatively it can be present pervasively across a taxonomic group [14, 15]. After *de novo* creation, a gene needs to become fixed in the population. The selective forces governing this process have been studied by examining the properties of orphan proteins. Intrinsic disorder, low complexity, subtelomeric location, high β-sheet preference as well as other features have been associated with orphan proteins [16, 17]. It has also been proposed that with age proteins (i) accumulate interactions, (ii) become more often essential and (iii) obtain lower *β*-strand content and higher stability [18]. Some aspects of these observations, such as the fact that orphans on average are short, are likely connected to a *de novo* creation mechanism. However, other features, including intrinsic disorder, are not obviously related to the gene genesis and could instead be the result of the selective pressure acting during fixation.

In yeast, we have earlier reported that the youngest proteins, i.e. the ones unique to *S. cerevisiae*, are less disordered than older proteins [7], while in *Drosophila* the opposite can be seen: the youngest proteins are more disordered than the ancient ones [19]. To the best of our knowledge the origin of this difference has not been explained. Could the selective forces be that disparate between two different eukaryotes? Or is it an artifact caused by limited genomic coverage? One difference between the two organisms is the content of Guanine and Cytosine (GC) nucleotides in the coding regions: *Saccharomyces cerevisiae* genomes are roughly 40% GC, while in *Drosophila melanogaster* the GC content is 53%.

To obtain a better understanding of the structural properties of orphan proteins, we determined the age of proteins in 187 eukaryotic genomes and studied a number of intrinsic properties, such as GC content, disorder, hydrophobicity, low complexity, and secondary structure. As expected we find that the most striking difference between young and old proteins is their difference in length. Further, intrinsic disorder and low complexity appear to be higher in orphans, albeit with a much smaller difference than for length, and these differences are not present in all species. The structural features in young proteins differ significantly depending on the GC content: low-GC orphans are much less disordered than high-GC orphans. In older proteins this relationship is much weaker, supporting a model where genes are created *de novo* starting from random DNA sequences, then their features gradually conform to those of ancient genes through adaptation.

## Materials and Methods

### Datasets

Protein data for 400 eukaryotic species were obtained from OrthoDB, release 8 [20], divided into 173 Metazoans and 227 Fungi, for a total of 4,562,743 protein sequences. This initial dataset was then filtered to a final size of 187 species, see below. For each species, a complete proteome was also downloaded from UniProt Knowledge Base [21].

### Age estimate

The ProteinHistorian software pipeline [13] is aimed at annotating proteins with phylogenetic ages. It requires a phylogenetic tree relating a group of species, and a protein family le representing the orthology relationships between proteins. The pipeline assigns each protein to an age group, depending on the species tree. Here, we used ProteinHistorian with default parameters, the NCBI phylogenetic tree [22], and protein orthology data obtained from OrthoDB. The OrthoDB method is based on all-against-all protein sequence comparisons using the Smith-Waterman algorithm and requiring a sequence alignment overlap of at least 30 amino acids across all members of an orthologous group. Therefore, the age group can be thought of as the level in the species tree on which a shared sequence of at least 30 amino acids first appeared, i.e. it assigns multi-domain proteins to the age of its oldest domains.

### Identification and definition of orphans

Proteins present in OrthoDB are only those with orthologs in at least one other species, i.e. proteins without orthologs (singletons) are not present in OrthoDB. Therefore, to obtain a set of candidate orphan proteins, the complete proteomes of all species were downloaded from Uniprot. Thereafter, BLAST was used to extract proteins not present in the OrthoDB dataset, obtaining 356,884 candidate orphan proteins. However, a large fraction of these proteins are not orphans but are missing from OrthoDB for other reasons, including that they were not present when the database was created or that they have undergone large domain rearrangements. We would assume that truly *de novo* created orphans do not contain domains found in other proteins. Therefore to ensure that we have a unique set of orphan proteins we filtered out proteins with hits in the Pfam-A database, by using hmmscan [23]. We believe that, due to the stringent criteria used here, the majority of this remaining set is constituted of *de novo* created proteins, and we refer to them as orphans throughout the rest of this paper. These proteins are specific to the species taxonomic level, i.e. we expect not to find them in other species in the dataset, even in the same genus. For *Saccharomyces cerevisiae*, that has 16 strains in the dataset, we also included the strain specific proteins into the orphan group.

Among the OrthoDB proteins, we defined genus orphans those that were assigned age = 1 (2 in the case of *S. cerevisiae*), while proteins having the maximum age according to ProteinHistorian were defined as ancient: these proteins are thought to be present in the common ancestor of all Fungi (taxon id = 4751) or all Metazoa (taxon id = 33208). Finally, proteins whose estimated age is between genus orphans and ancient were defined as intermediate.

Taxonomic genera represented by a single species in the dataset have by definition no genus orphans; for this reason, we selected for our final dataset only the 187 species that have at least one other species within the same genus. The final dataset amounts to 1,782,675 proteins distributed across 187 species; 0.8% of them are defined orphans and 0.6% as genus orphans, 15% are intermediate and the remaining 84% are ancient.

One problem that exists using the NCBI phylogenetic tree is the presence of many polytomic branches, especially at the genus level, because ProteinHistorian cannot distinguish between proteins being specific to that species and proteins shared among the entire group. To solve this, we forced no polytomy on the terminal branches: multifurcating nodes were converted to a randomly bifurcated topology, transforming the NCBI tree to a fully binary structure. While a binary tree is needed for ProteinHistorian, its algorithm assumes that a protein gain is much more rare than a loss; this means that the most recent common ancestor of a protein will be at the topmost intersection of a group of species. Thus, randomly converting multifurcations to bifurcation might likely underestimate the number of genus-specific orphans, but have no effect on species-specific orphans.

Clades affected by the conversion from multi-to bi-furcating branches include *Caenorhabditis* (5 species), *Drosophila* (5 species), *Anopheles* (5 species), *Candida* (5 species), *Saccharomyces* (14 strains), *Aspergillus* (5 species) and *Trichopython* (5 species). The taxonomic tree comprising the final set of 187 species is presented in Fig. S1.

### Assigning GC content

We could map 1,357,518 out of 1,782,675 proteins (~76% of the dataset) to their ENA identifiers. This mapping was used to download the Coding Sequence (CDS) data from ENA (*https://www.ebi.ac.uk/ena/*); the GC content was then calculated for each mapped gene individually.

### Gene ontology annotation

Evidence for functionality of the proteins was estimated using annotated Gene Ontology (GO) terms. Using the Uniprot KnowledgeBase mapping data *ftp://ftp.uniprot.org/pub/databases/uniprot/currentrelease/knowledgebase/idmapping/idma13p1ping.dat.gz* we assigned UniprotKB identifiers to 894,831 out of 1,752,675 proteins (51%). These were then annotated with three terms, one for each main GO category: Molecular Function, Biological Process and Cellular Component. All GO terms are associated with evidence codes; a subset of these codes (‘EXP’, ‘IDA’, ‘IPI’, ‘IMP’, ‘IGI’ or ‘IEP’) represents experimentally validated functional annotations. If any of these codes is present we mark the corresponding protein as experimentally characterized.

### Predicted properties of proteins

Intrinsic disorder content was predicted for all the proteins by using several disorder predictors; short and long disorder predictions by IUPred [24], three type of predictions (REM-465, Hotloops and Coils) by DisEMBL [25] and GlobPlot [26]. In the main figures we only report the prediction by IUPred long; the others are found in the supplementary material (Fig. S3 to S8). It is worth mentioning that these predictors operate with different definitions of disorder, so a consensus should not be expected.

We used SCAMPI [27] to predict the fraction of transmembrane residues in a protein. The fraction of low-complexity residues is predicted using SEG [28]. PSIPRED [29] was used to predict the secondary structure of all the proteins in the dataset, using only a single sequence and not a profile. This reduces the accuracy but the overall frequencies should not be changed significantly. We annotated each protein with the fraction of residues predicted to be in each type of secondary structure (α-helix, β-strand, coil).

### Propensity scales

TOP-IDP [30] is a measure of the disorder-promoting propensity of a single amino acid. For each protein, the average propensity was calculated by averaging the TOP-IDP values of all its residues. Similarly the hydrophobicity of each protein was expressed as the average hydrophobicity using the biological hydrophobicity scale [31]. Finally, we computed the propensity of each amino acid to be in a secondary structure (helix, sheet, coil, turn) in the same manner by using secondary structure propensity scales [32].

### Statistical significance of the results

In order to test the statistical significance of the results, a number of tests were performed. Rank-sum tests between all possible pairs of age groups were performed for the entire dataset and for each studied property. Due to the large number of samples the p-values from these tests are always smaller than 10^−141^ even when the absolute difference in numbers is minuscule.

To study the difference between young and old proteins on a global level, we performed a rank-sum test for orphan versus ancient proteins within each species. To exclude small variations we only considered the species where the p-value of this test was <0.01.

To determine the relationship between a property and GC we studied the slopes for proteins of different age. If the p-value of a linear regression test is <0.01, the corresponding property is considered significantly correlated with GC.

### Random proteins at different GC contents

To test whether the studied intrinsic properties, as well as the frequency of any given amino acid, were solely dependent on GC content, we used a set of 21,000 random ORFs, generated as follows: at each GC content ranging from 20 to 90%, in steps of 1%, a set of 400 ORFs (equally divided into 300, 900, 1,500 and 2,100 bp long) was generated so that its content of GC was fixed. The ORFs were generated by randomly selecting codons among the 61 non-stop codons. The probability to select one codon given a GC content of GC_*freq*_ is set accordingly:

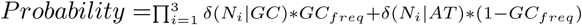

Where *N*_*i*_ is the nucleotide of the codon in position *i* and *δ*(*N|GC* is equal to 1 if the nucleotide N is guanine or cytosine and zero otherwise, etc. Finally, start and stop codons were added. These ORFs were then translated to polypeptides, and all their intrinsic properties, as well as the frequencies of their amino acid were computed, as described above.

## Results

**Figure 1.**
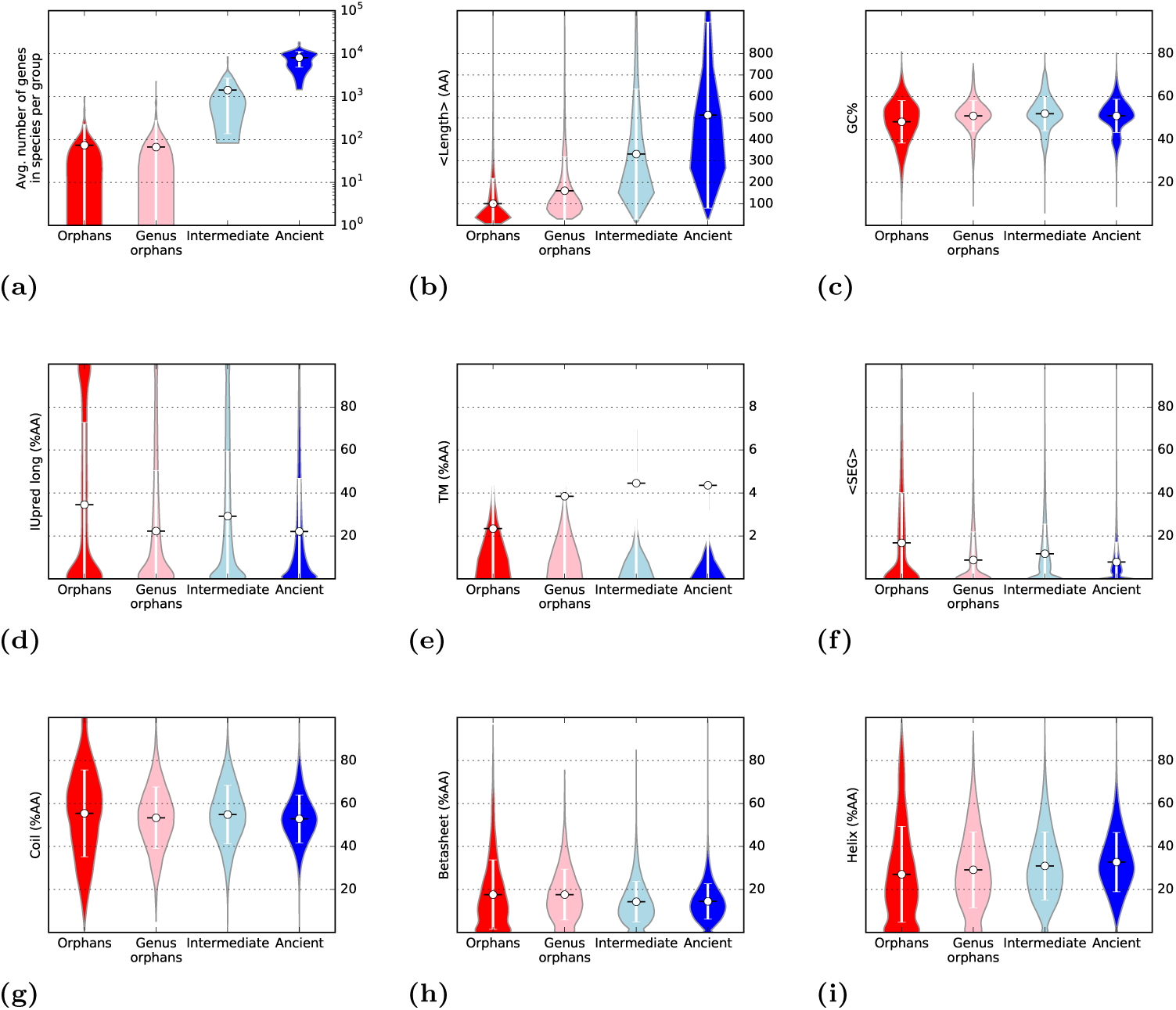
Overview of the proteins assigned to the four age groups: (a) the fraction of proteins belonging to each age group, (b) the average length, in amino acids, (c) the average GC content of the genes, (d) intrinsic disorder predicted by IUpred (long), (e) percentage of transmembrane residues, (f) percentage of residues in low-complexity regions, percentage of residues predicted to be in (g) a coil, (h) a *β*-sheet and (i) in a helix. The difference between orphans and ancient is statistically significant for all the considered properties: the p-value of a rank-sum test is always < 10^−141^.

The assignment of age to all proteins is based on the ProteinHistorian pipeline [13]. In the youngest, orphan group, only proteins that are (a) not present in any other genome and (b) that do not share any Pfam-A domain with any other eukaryotic protein are present. In the next group, genus orphans, only proteins that are unique to a genus are included. Proteins having the maximum age according to ProteinHisorian are labeled as ancient; the rest is classified as intermediate.

A summary of the age assignment and subsequent grouping into orphans, genus orphans, intermediate and ancient is shown for each species in Table S1. Orphans and genus orphans constitute each less than 1% of the dataset; intermediate proteins are ~15% and ancient proteins ~84%

These results show that for most genomes we do a conservative estimate of the number of orphans and find fewer orphans than in earlier studies. For instance, in *Saccharomyces cerevisiae* we identified 16 orphans and 5 genus orphans, out of 6466 proteins. As a comparison, in our earlier study we reported 157 species-specific and 125 genus-specific orphans [7] while Vidal and co-workers reported 143 species-specific (ORFs_1_) and 609 genus-specific (ORFs_2—4_) proteins [33]. Similarly, in *Drosophila pseudoobscura* we could identify only 6 orphan proteins, in comparison to the 228 reported previously [8]. This shows that the exact identification of which proteins are *de novo* created remains difficult and depends on the genomes included in the study.

However, our primarily aim for this study is not to estimate the exact number of orphans, but to examine properties of proteins of different ages. Therefore, we do believe that our conservative estimate is useful to enhance the fraction of *de novo* created proteins in the youngest groups.

### Functional annotations

**Table 1.** For the four age groups, the fraction of proteins being annotated in Gene Ontology (GO) is shown. In parentheses is the fraction of proteins having an experimentally validated annotation (i.e. a GO evidence code equal to ’EXP’, ’IDA’, ’IPI’, ’IMP’, ’IGI’ or ’IEP’).

| Group | GO Process | GO Function | GO Component |
| --- | --- | --- | --- |
| Ancient | 25.7% (0.8%) | 26.7% (0.5%) | 22.4% (0.8%) |
| Intermediate | 9.2% (0.5%) | 10.6% (0.2%) | 12.6% (0.4%) |
| Genus Orphans | 7.4% (0.0%) | 7.4% (0.0%) | 14.1% (0.0%) |
| Orphans | 3.3% (0.3%) | 3.5% (0.1%) | 9.4% (0.3%) |

Next we set out to estimate the functional evidence for our set of proteins; for this we explore their Gene Ontology (GO) annotation. For each main GO category (process, function and component), we computed the fraction of proteins being annotated with at least one GO term in UniProt. In addition we calculated the fraction of proteins having at least one experimentally verified GO annotation, Table 1.

The fraction of annotated proteins increases steadily with age, from ~3-9% in orphans to ~25% in ancient, Table 1. This is expected, as older proteins have more regulatory, protein-protein, and genetic interactions [18]. However, the fraction of proteins with experimental functional evidence is small (<1% of protein) irrespectively of age. This shows that there exists at least a fraction of proteins of any age that is functionally characterized, but it is difficult to exactly determine how substantial it is.

### In most genomes orphan proteins are more disordered

The average length of the proteins increases by age, see Fig 1b. The average length is 100 amino acids in orphans, 150 in genus orphans, 300 for intermediate and 500 for ancient proteins. It can be noted that in the vast majority of the genomes the difference in length is significant between orphan and ancient proteins, Table 2. This highlights the well-established fact that eukaryotic proteins expand during evolution: the expansion can occur by several mechanisms, including domain-fusions [34], additional secondary structure elements [35] and expansion within intrinsically disordered regions [16].

GC content on the other hand does not appear to change by age, see Fig 1c. There is approximately the same number of genomes where a statistically significant increase or decrease exist, Table 2.

**Table 2.** For the 187 considered species, the number of species in which a property is significantly higher (increasing) or significantly lower (decreasing) in orphans compared to ancient proteins is shown.

| Property | # > in orphans | # > in ancient |
| --- | --- | --- |
| General properties |  |  |
| Length (AA) | 0 | 135 |
| GC% | 23 | 33 |
| Disorder predictors |  |  |
| IUPred long (%AA) | 64 | 12 |
| IUPred short (%AA) | 103 | 2 |
| DisEMBL Coils (%AA) | 96 | 3 |
| DisEMBL REM465 (%AA) | 125 | 0 |
| DisEMBL Hotloops (%AA) | 111 | 1 |
| GlobPlot (%AA) | 69 | 4 |
| Secondary structure predictions |  |  |
| SEG (%AA) | 45 | 3 |
| Coil (%AA) | 40 | 9 |
| Helix (%AA) | 2 | 51 |
| Sheet (%AA) | 13 | 13 |
| TM (%AA) | 1 | 17 |
| Propensity scales |  |  |
| Alpha propensity | 29 | 9 |
| Beta propensity | 20 | 18 |
| Coil propensity | 14 | 33 |
| Turn propensity | 11 | 35 |
| Hydrophobicity | 22 | 28 |
| TOP-IDP | 50 | 17 |

Next, we compared predicted structural properties of all proteins, see Fig. 1d-l, S2 and Table 2.

The amount of predicted disorder residues ranges between 20% and 40%, depending on the prediction method. For most disorder predictors the fraction of disordered residues is higher in orphans than in ancient proteins. However, there exists about a handful of genomes where the opposite trend is observed: supporting earlier observations, orphans are significantly more ordered in *Candida albicans* according to 5 out of 6 methods, in *Saccharomyces cerevisiae s288c* for 4 methods and in *Fusarium pseudograminearum* for 3. An interesting case is that of *Drosophila pseudoobscura*, that appears to have more ordered orphans according to IUPred long, contrary to all others *Drosphila* species.

The fraction of transmembrane residues is on average ~2% in orphan proteins, with an increasing trend towards ancient (4%). Similarly the amount of helical residues increase slightly with age, while the fraction of low complexity residues decrease by age. For all these structural predictions the changes are quite small and there are genomes with significant increases and decreases for all measures.

### Orphan proteins are more disordered in yeast but less in *Drosophila*

Above, we noted that on average orphan proteins are more disordered. However, we also noted that in a handful of genomes a statistically significant opposite trend could be observed. To investigate this further we studied the amount of predicted disorder in each genome separately. When studying intrinsic disorder, orphans and genus orphans of *S. cerevisiae* appear remarkably ordered (~3% of the amino acids) as shown before [7] see Fig. 2a and b. The closely related species *Candida albicans* shows a similar trend; see Fig. 2c. Results from additional disorder predictors are presented in Fig. S4-8 and agree well with these observations.

In contrast, but also consistent with earlier studies [36], orphans and genus orphans in most *Drosophila* genomes are more disordered than ancient proteins, see Fig. 2d and e. In the worm *C. elegans* (Fig. 2f) orphan proteins appear to be consistently more disordered than progressively older ones; this is true across all the considered *Caenorhabditis* species. These results are consistent in other predictors, see Fig. S4-8 d, e and f.

In general, it is apparent that in most organisms orphans are more disordered than ancient proteins, while in yeast the opposite appears to be the case. What could possibly explain this difference? One possibility is that the more complex regulations in animals require more disordered residues in comparison with yeast. But the average disorder content is similar in all eukaryotic species, contradicting this idea.

We noted that yeasts are among the genomes with lowest GC content (~40% in *S.cerevisiae*, 35% in *C. glabrata*). Therefore, we decided to examine the properties of proteins from different age groups in respect to their GC content.

**Figure 2.**
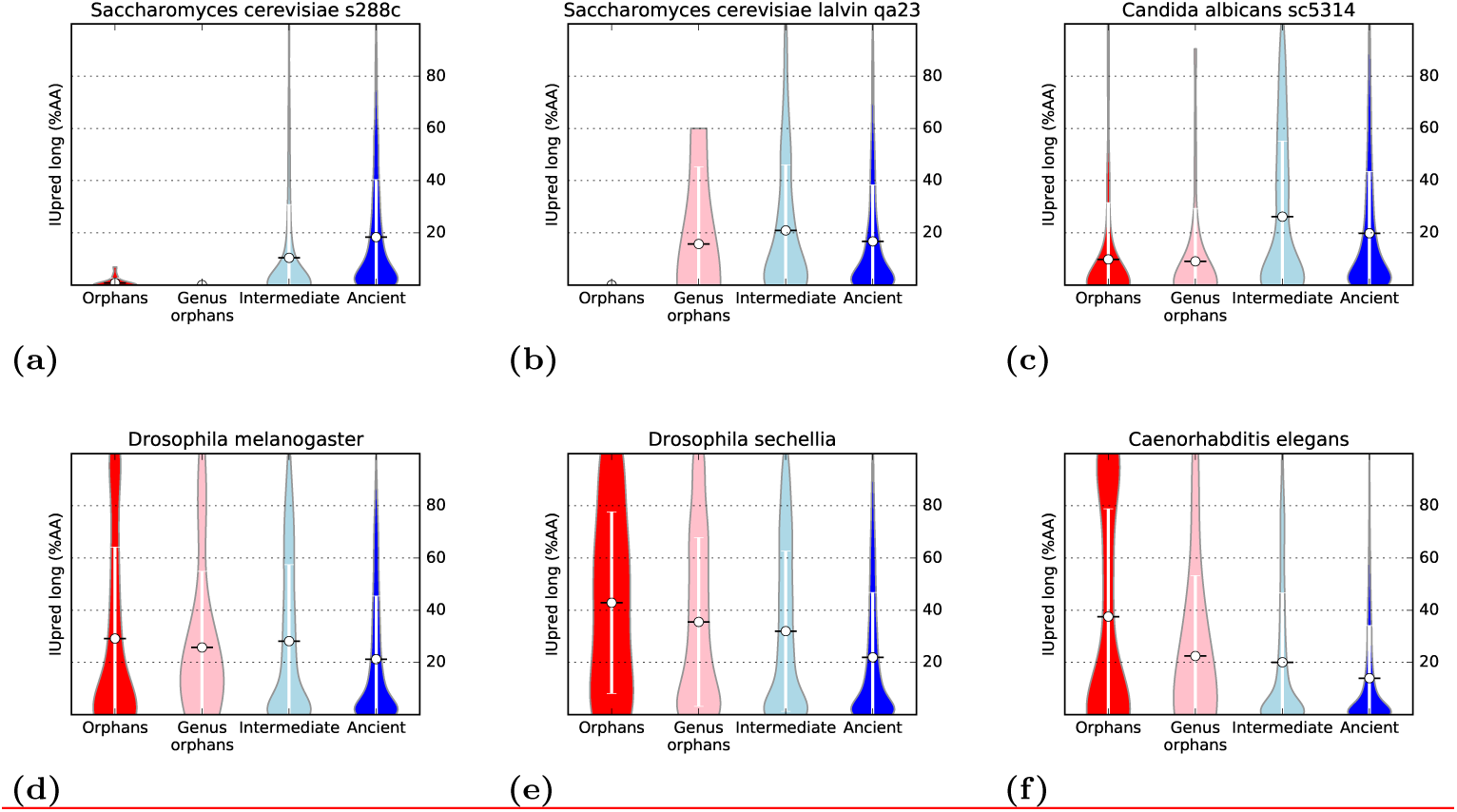
For six selected species ((a,b) two strains of *S. cerevisiae*, (c) *C. Albicans*, (d) *D. melanogaster*, (e) *D. sechellia* and (f) *C. elegans*), intrinsic disorder (% of amino acid predicted as disordered by IUpred long) is shown as violin plots for proteins in the different age groups.

### Orphans are more disordered in high-GC genomes

To identify the origin of the different properties of orphan and ancient proteins in different organisms we studied the distribution of different structural properties, including low complexity, fraction of transmembrane residues, secondary structure frequency and intrinsic disorder) for all genomes against GC of the genomes, see Fig. 3.

With the exception of *β*-sheet frequency, the difference between orphans and ancient proteins for all the considered properties is statistically significant: the p-value of a rank-sum test (a non-parametric equivalent of the t-test) is always < 10^-11^.

**Figure 3.**
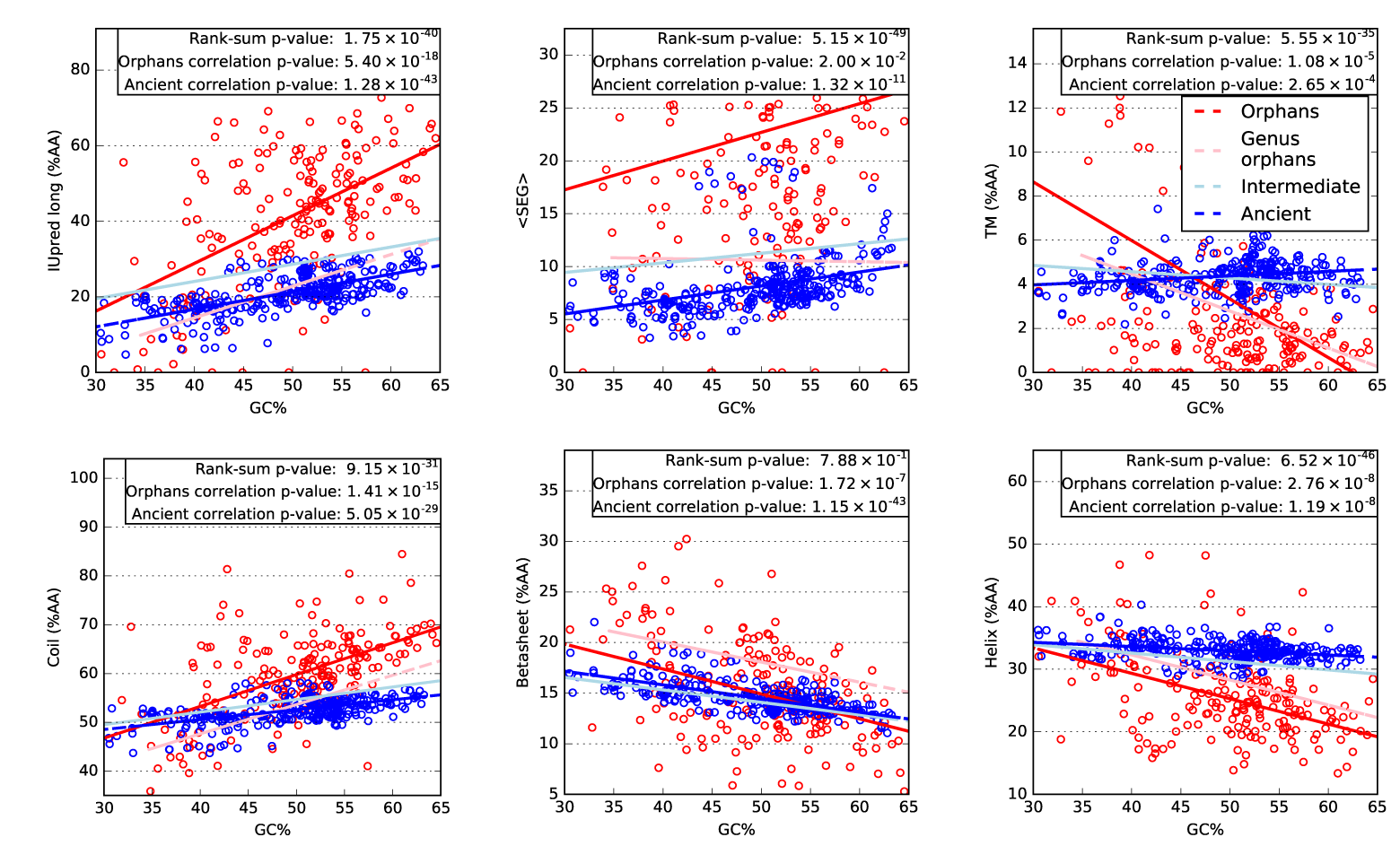
Structural properties of proteins of different ages plotted against the GC content of the genome (coding regions). For clarity only the ancient (blue) and orphan (red) proteins are shown individually, but the linear fitted lines for genus orphans (pink line) and intermediate ones (light blue) are also shown. In the text box three values are presented: rank-sum p-value = p-value of a rank-sum test of orphans versus ancient (only the property on y axis is considered); correlation p-values = p-value of a linear regression test for orphan and ancient.

For proteins of all ages, low complexity (SEG) and predicted coil frequency increase with GC, while transmembrane, helix and sheet frequency decrease.

Notable is that intrinsic disorder shows a clear, directly proportional dependency on GC: higher GC corresponds to more disorder. At the extreme (over 60% GC), more than 50% of the residues in orphan proteins are predicted to be disordered, while for ancient proteins the disorder fraction is about 30%. At low GC (below 40%) the fraction of disordered residues is lower and similar in ancient and orphan proteins (15-20%).

Further, the dependency of GC is clearly stronger for younger proteins, indicating that it is related to the creation of the protein and then gradually lost during evolution.

To assess the significance of this dependency, we performed a linear regression test for each age group. The p-values of such test is presented for orphans and ancient in the boxes of Fig. 3. All the properties, with the exception of low complexity, show a p-value <0.01, indicating that they are significantly correlated with GC in both orphan and ancient proteins.

The GC is not constant over a genome. In complex eukaryotic organisms, the global GC content is heavily determined by the GC composition of isochores: these regions of uniform GC form a mosaic in the genomes of many complex eukaryotes, and their maintenance is likely the result of natural selection [37]. In general coding regions have higher GC than noncoding regions [38,39]. Further, there are also variation in GC between different regions of a genome, so when a noncoding region is turned into a gene the local GC will decide the amino acid content of the protein.

Therefore, it might be more relevant to study the GC of each protein individually.

### A strong relationship between GC and structural properties of orphans

In Fig. 4 we show the dependency of structural properties on GC content for individual proteins. In addition, structural properties of a set of proteins generated randomly at all GC levels are shown. Orphans and genus orphans, as well as random proteins, show a definite dependency on GC. In contrast, ancient and intermediate proteins are only loosely dependent on GC.

In general there is a resemblance between orphans and randomly generated proteins. However, when studying Fig. 4 in more detail a few notable differences between them can be observed: orphans are more disordered and contain more low complexity regions but fewer sheets, independently of the GC level.

It should be recalled that what we describe above is based on *predicted* structural features that are an indirect reflection of the protein sequence. If a certain group of proteins is predicted to be more disordered, or contain more sheets, it is quite likely a consequence of changes in amino acid frequencies.

**Figure 4.**
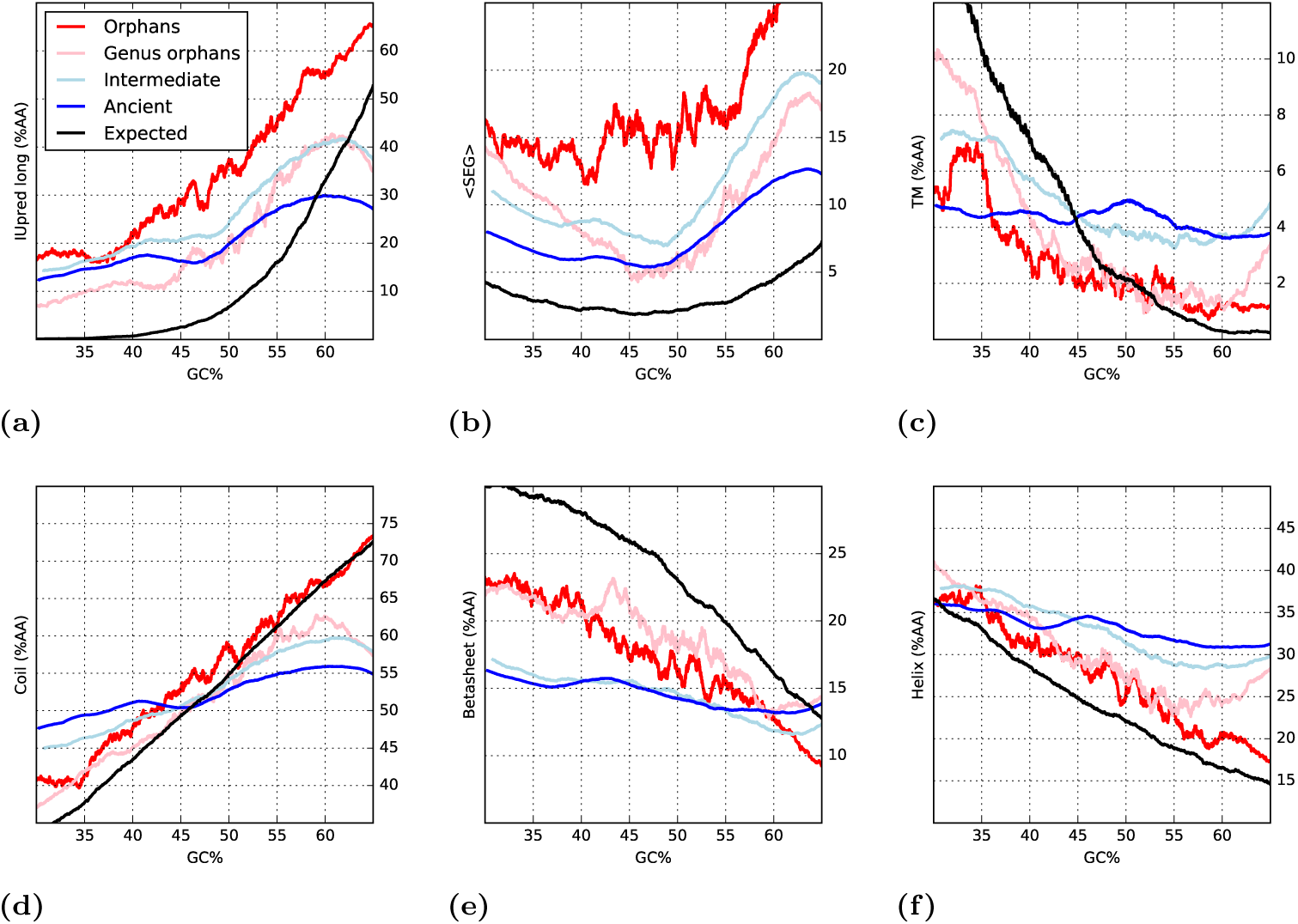
Running averages of predicted structural properties against GC content: (a) disorder, predicted by IUpred (long); (b) low complexity, predicted by SEG; (c) percentage of transmembrane residues predicted by Scampi; (c,d,e) percentage of residues in secondary structure of type, respectively, coil, beta sheet and alpha helix. For each property, colored lines represent proteins of different age: orphans (red), genus orphans (pink), intermediate (light blue) and ancient (blue). The black lines represent randomly generated proteins at different GC frequencies.

### Property scales

Next, we studied the proteins using six different amino acid propensity scales. The difference between the scales and predicted features is that scales describe general properties, and are directly calculated from amino acid frequencies, while predicted properties can also include other features. For disorder we use the TOP-IDP scale [30], for hydrophobicity we use the biological hydrophobicity scale [31], while sheet, turn, coil and helix propensities are analyzed using the structure-based conformational preferences scales [32].

**Figure 5.**
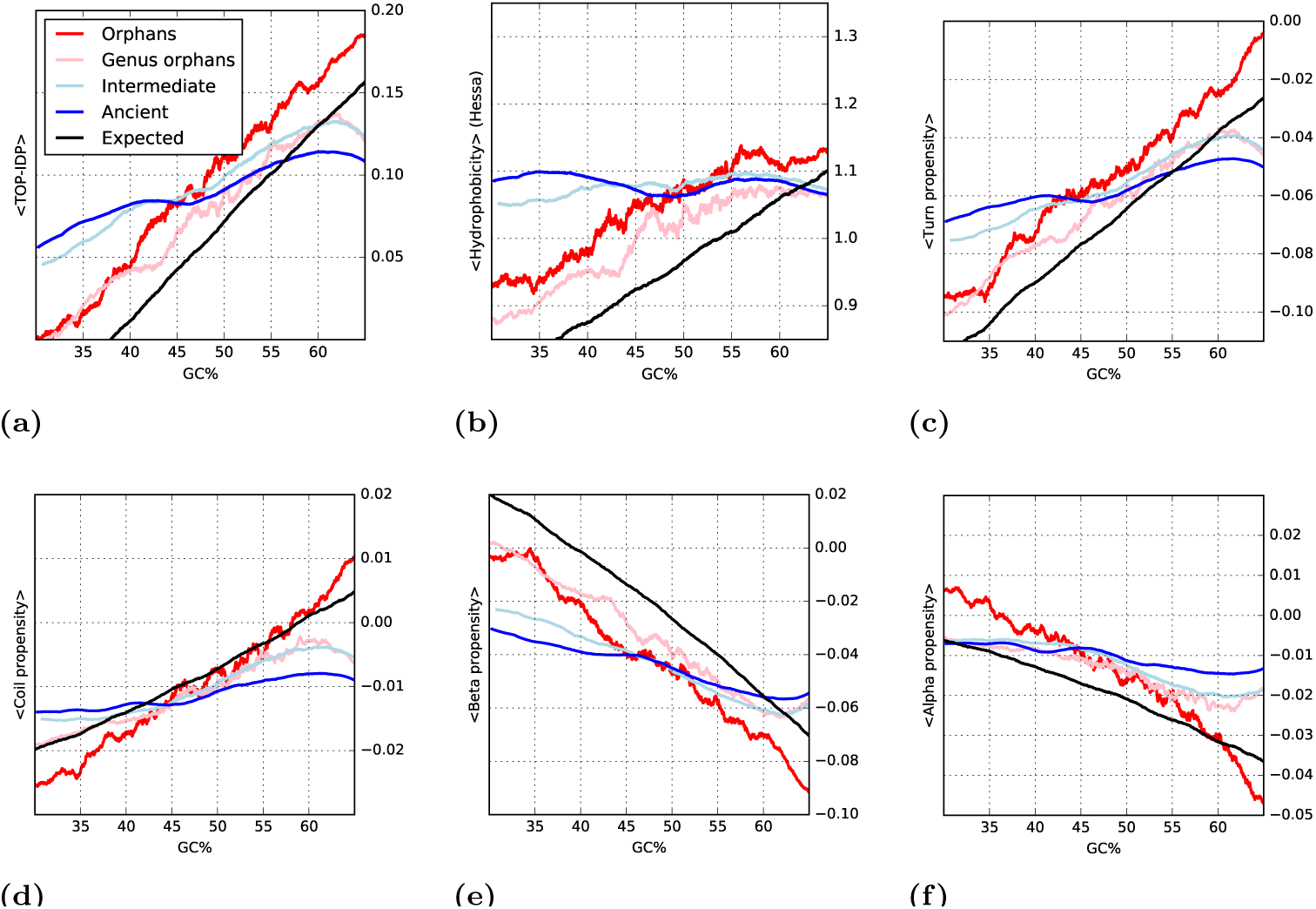
Running averages of structural properties computed from amino acid scales against GC content: (a) Intrinsic Disorder Propensity (TOP-IDP); (b) hydrophobicity (Hessa scale); (b, c, d, e) average propensity for secondary structure of type, respectively, turn, coil, beta sheet and alpha helix. For each property, colored lines represent proteins of different age: orphans (red), genus orphans (pink), intermediate (light blue) and ancient (blue). The black lines represent randomly generated proteins at different GC frequencies.

In agreement with the predicted values; the average properties in the four age groups of proteins are overlapping, see Fig. S2. However, when taking the GC content into account the properties of the younger proteins show a strong correlation with GC, see Fig. 5. To a very large degree the properties of orphan proteins follow what would be expected for random proteins (black line). However, regardless of GC, orphan proteins are more disordered and hydrophobic, have slightly higher turn and helical propensities, and also lower sheet propensities than random.

Interestingly, the propensities of the two groups of older proteins also change by GC; however, this dependency is less pronounced than for orphan or random proteins. The difference seen between orphan and ancient proteins indicates that, given evolutionary time, the selective pressure to change the GC level is weaker than the selective pressure to change amino acid frequencies.

**Figure 6.**
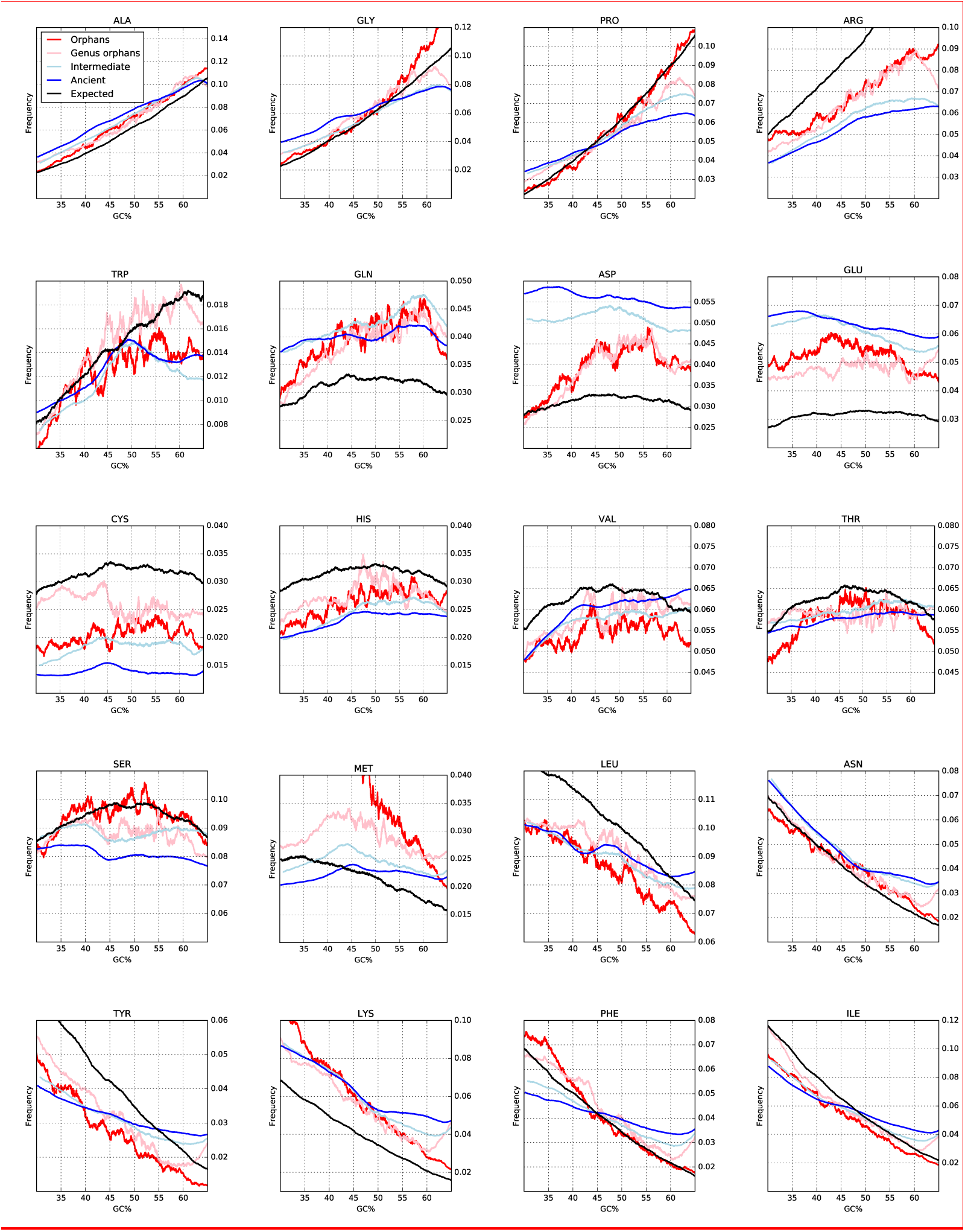
The relationship of each amino acid frequency with the GC content and age of the protein. A black line represents the expected values. The amino acids are sorted by the GC content in their codons.

## Discussion

GC content affects the codon usage in the genome [40] and it has been argued that GC might be the sole responsible for codon bias [41]. The difference in codon usage causes differences in amino acid frequencies in such a way that some amino acids are more frequent in higher GC for random protein sequences. Also, a preference for some amino acids might cause a change in GC just by a higher frequency of certain codons. In younger proteins the correlation between amino acid frequency and GC is stronger than in older proteins, see Fig. 5. This indicates that the selective pressure to change amino acids in a protein is stronger than the one to change the GC content. At low GC ancient proteins are more disordered than expected for random sequence while at high GC they are less.

**Figure 7.**
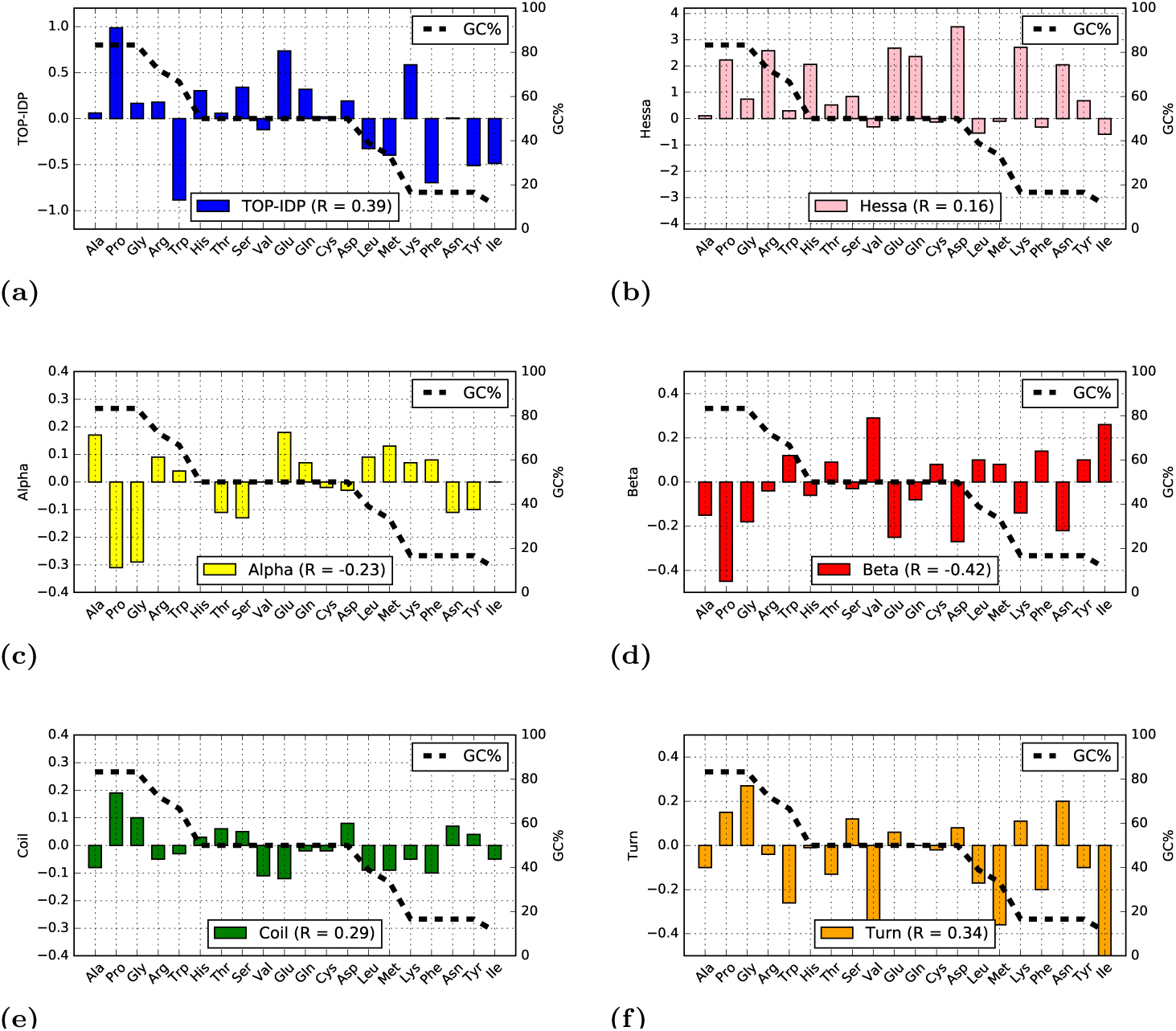
The fraction of GC in all codons encoding an amino acid is plotted as a dotted line and the values for the different propensity scales as filled bars: (a) TOP-IDP, (b) Hessa transmembrane scale, and (c-f) Koehl secondary structure preference scale. For each scale the Pearson (R) correlation with GC is also shown.

### The influence of GC on amino acid preferences

How do changes in GC content affect proteins? In a random DNA sequence, the frequency of different codons changes depending on GC, and this, in turn, affects the resulting amino acid frequencies. To study the effect of GC content on amino acid frequency we examined the frequency of all 20 amino acids in proteins of different age.

In Fig. 6, the expected and observed amino acid frequencies at different GC contents are explored. For most amino acids the observed frequencies are surprisingly well correlated with what is expected from GC alone. However, a few notable exceptions exist:

- For Pro, Arg, Trp, Tyr, Phe and Ile, the frequencies in orphan proteins resemble the random proteins and are strongly dependent on GC content, while the frequencies in ancient proteins are much less dependent on GC content. This suggests that there is a selective pressure to gradually adjust the frequencies of these amino acids to an optimal level.
- Asn and Ala change in both orphans and ancient proteins with GC frequency, indicating that the selective pressure to change the frequency of these amino acids is lower.
- Glu, Gln and Asp are more frequent than expected, at any GC level. Their frequency in orphans is intermediate to what is expected by chance and what is found in ancient proteins, suggesting a gradual increase during evolution. These amino acids are coded by only two codons, i.e. there exists a selective pressure to increase their frequency to a higher level than the 3.3% expected by chance.
- Finally, Cys and His are less frequent, independently of GC content, in real proteins than in random ones, indicating their special roles in protein function and folding as well as their rareness.

In Fig. 7 and Table S2 the GC content of the codons of each amino acid is compared with the propensity of that amino acid to be in a certain structural region. Three amino acids, Ala, Gly and Pro are “high GC” amino acids, i.e. they have more than 80% GC in their codons, while five amino acids, Lys, Phe, Asn, Tyr and Ile, are “low GC codons”, with less than 20% GC in their codons.

All three “high GC” amino acids are disorder promoting (high TOP-IDP), and four out of five “low GC” amino acids are order-promoting (low TOP-IDP) residues. Therefore at high GC content, codons coding for hydrophilic, disorder-promoting amino acid are prevalent. Genes low in GC tend to contain codons for hydrophobic amino acids, associated with order.

All scales correlate with the GC frequencies with coefficients ranging from -0.42 to 0.39. The strongest correlations are found with β-sheet propensity (-0.42) and TOP-IDP (0.39) and the weakest with hydrophobicity (0.16).

## Conclusions

We have studied the properties of proteins and their age in a large set of eukaryotic genomes. As shown before, orphan proteins are shorter than ancient proteins, but, surprisingly, we do find that on average for other structural features the young and old proteins are rather similar. However, we also observe that the properties of youngest proteins vary significantly with the GC content. At high GC the youngest proteins become more disordered and contain less secondary structure elements, while at low GC the reverse is observed. We show that these properties can be explained by changes in amino acid frequencies caused by the different amount of GC in different codons. The influence of this can be seen in the frequency of the amino acids that have a high or low fraction of GC in their codons, such as Pro.

In a random sequence, Pro only represents less than 5% of the amino acids at 40% GC, but 10% at 60% GC. This actually agrees well with what is observed in orphan proteins: 5% at 40% GC vs. 9% at 60% GC, see Fig. 6. Similar changes in frequencies can be observed for several amino acids.

On average, young proteins are more disordered than ancient proteins, but this property is strongly related to the GC content. In a low-GC genome the disorder content of an orphan protein is ∼30% while in a high-GC genome it is over 50%, see Fig. 3.

Here we show that GC content of a genome strongly affects the amino acid distribution in *de novo* created proteins. It appears as if *de novo* created proteins that become fixed in the population are very similar to random proteins given a certain GC content. Codons coding for disorder-promoting residues are on average richer in GC, explaining the earlier contrasting observations between the low disorder among orphans in yeast (a low GC organism) and the high disorder among orphans in *Drosophila* (a high GC organism).

Finally, it can be observed that older proteins show a lower dependency of their structural properties on GC, but have a GC content similar to the one of orphans. This can lead to the speculation that selective pressure acts less on GC levels and more on structural features of proteins.

## Supporting Information

**Table S1.**
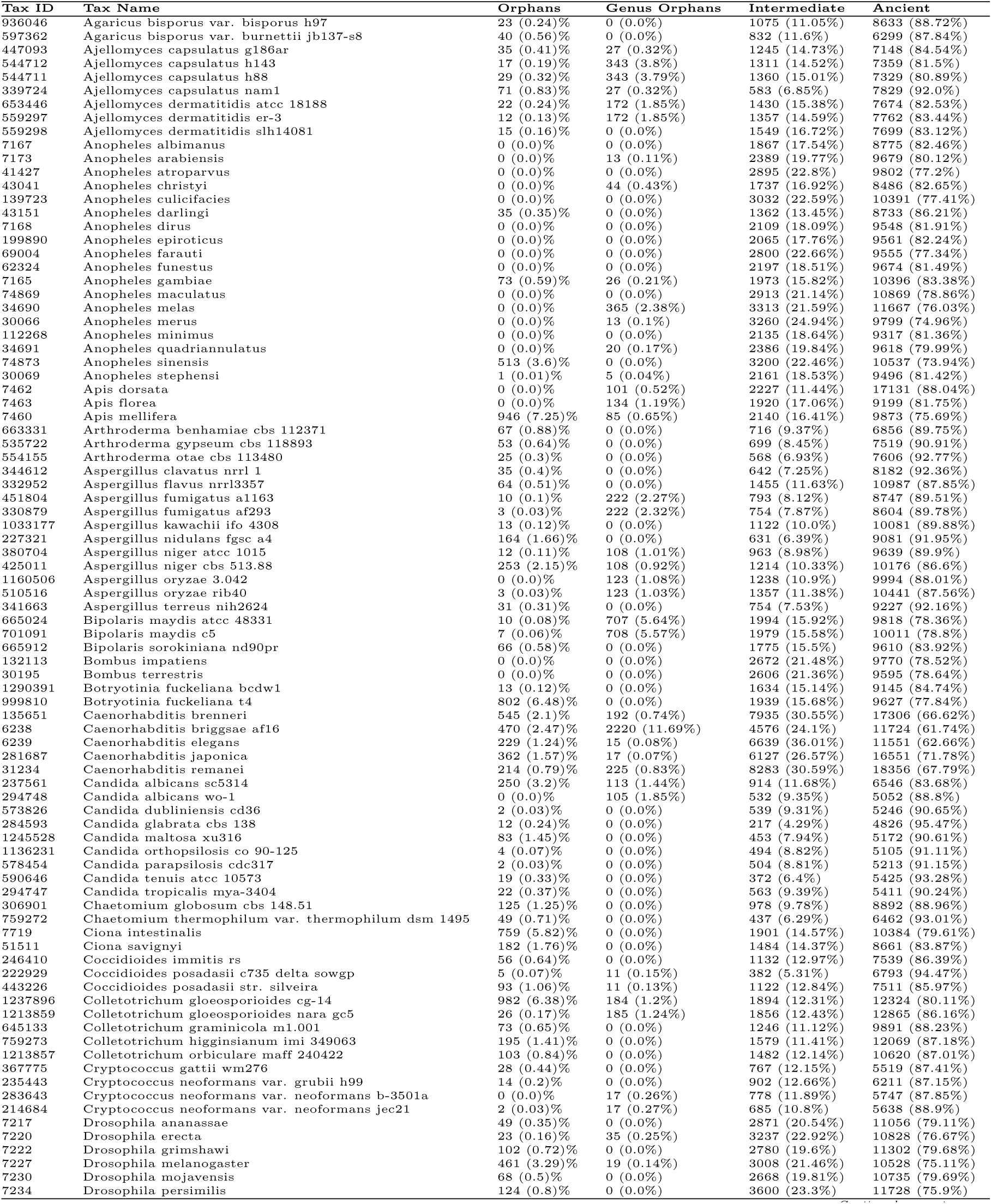
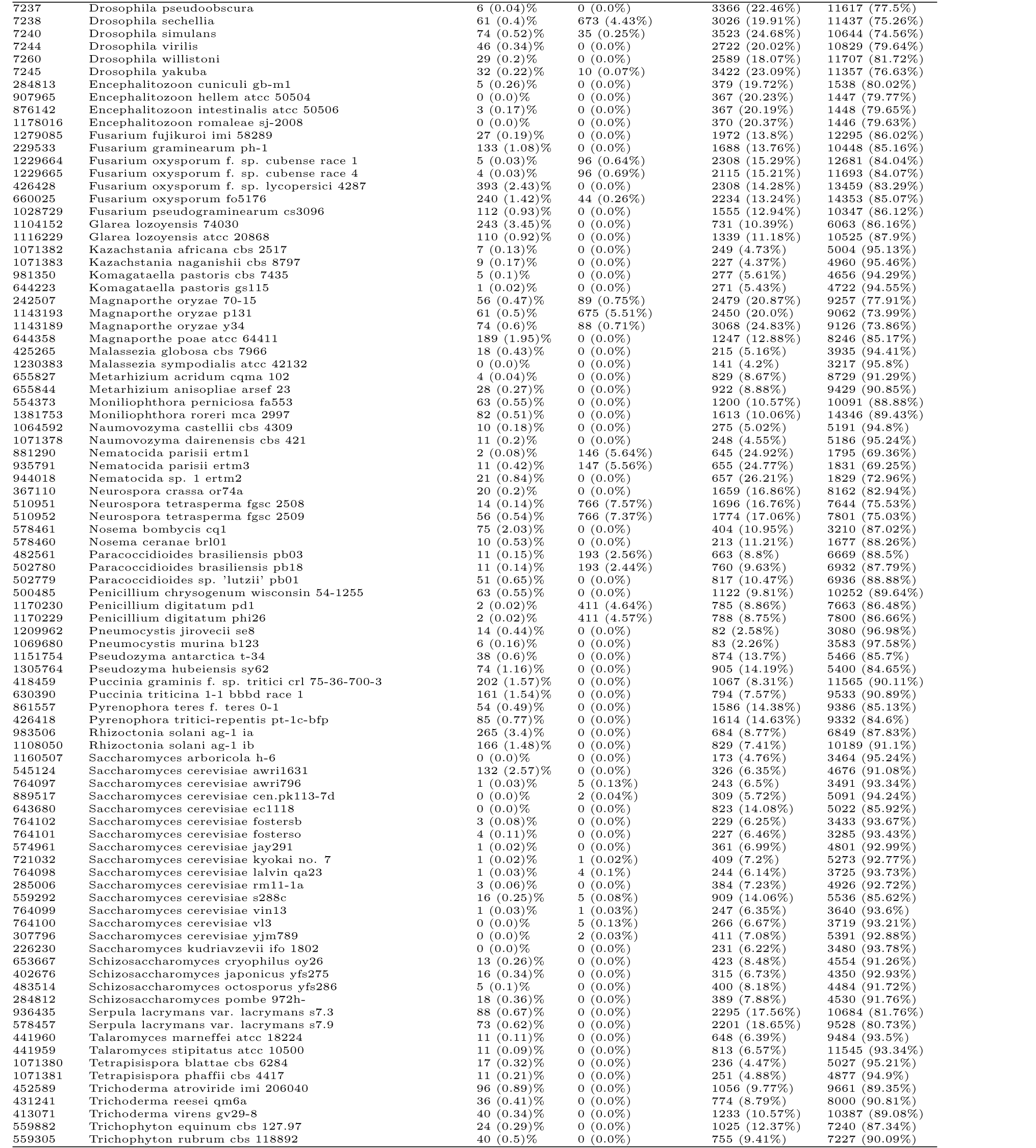
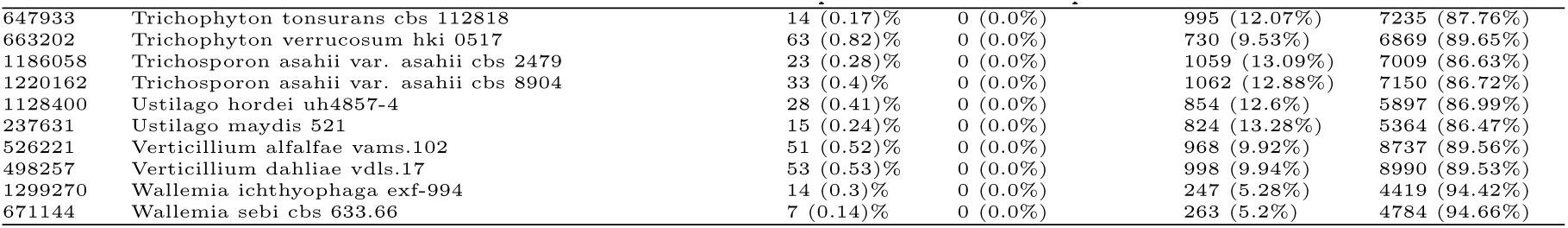
Summary of the numbers (with percentages in parentheses) of orphans, genus orphans, intermediate and ancient proteins for each of the 187 considered eukaryotic species.

**Table S2.** For each amino acid we show the GC content of its codons, its value in the TOP-IDP scale, its hydrophobicity (Hessa scale), as well as its value in the secondary structure scales for Alpha, Beta, Coil and Turn propensity.

| AA | GC% | TOP-IDP | Hessa | Alpha | Beta | Coil | Turn |
| --- | --- | --- | --- | --- | --- | --- | --- |
| Ala | 0.83 | 0.06 | 0.11 | 0.17 | -0.15 | -0.08 | -0.10 |
| Arg | 0.72 | 0.18 | 2.58 | 0.09 | -0.04 | -0.05 | -0.04 |
| Asn | 0.17 | 0.01 | 2.05 | -0.11 | -0.22 | 0.07 | 0.20 |
| Asp | 0.50 | 0.19 | 3.49 | -0.03 | -0.27 | 0.08 | 0.08 |
| Cys | 0.50 | 0.02 | -0.13 | -0.02 | 0.08 | -0.02 | -0.02 |
| Gln | 0.50 | 0.32 | 2.36 | 0.07 | -0.08 | -0.02 | 0.00 |
| Glu | 0.50 | 0.74 | 2.68 | 0.18 | -0.25 | -0.12 | 0.06 |
| Gly | 0.83 | 0.17 | 0.74 | -0.29 | -0.18 | 0.10 | 0.27 |
| His | 0.50 | 0.30 | 2.06 | 0.00 | -0.06 | 0.03 | -0.01 |
| Ile | 0.11 | -0.49 | -0.60 | 0.00 | 0.26 | -0.05 | -0.62 |
| Leu | 0.39 | -0.33 | -0.55 | 0.09 | 0.10 | -0.09 | -0.17 |
| Lys | 0.17 | 0.59 | 2.71 | 0.07 | -0.14 | -0.05 | 0.11 |
| Met | 0.33 | -0.40 | -0.10 | 0.13 | 0.08 | -0.09 | -0.36 |
| Phe | 0.17 | -0.70 | -0.32 | 0.08 | 0.14 | -0.10 | -0.20 |
| Pro | 0.83 | 0.99 | 2.23 | -0.31 | -0.45 | 0.19 | 0.15 |
| Ser | 0.50 | 0.34 | 0.84 | -0.13 | -0.03 | 0.05 | 0.12 |
| Thr | 0.50 | 0.06 | 0.52 | -0.11 | 0.09 | 0.06 | -0.13 |
| Trp | 0.67 | -0.88 | 0.30 | 0.04 | 0.12 | -0.03 | -0.26 |
| Tyr | 0.17 | -0.51 | 0.68 | -0.10 | 0.10 | 0.04 | -0.10 |
| Val | 0.50 | -0.12 | -0.31 | 0.00 | 0.29 | -0.11 | -0.41 |

**Figure S1.**
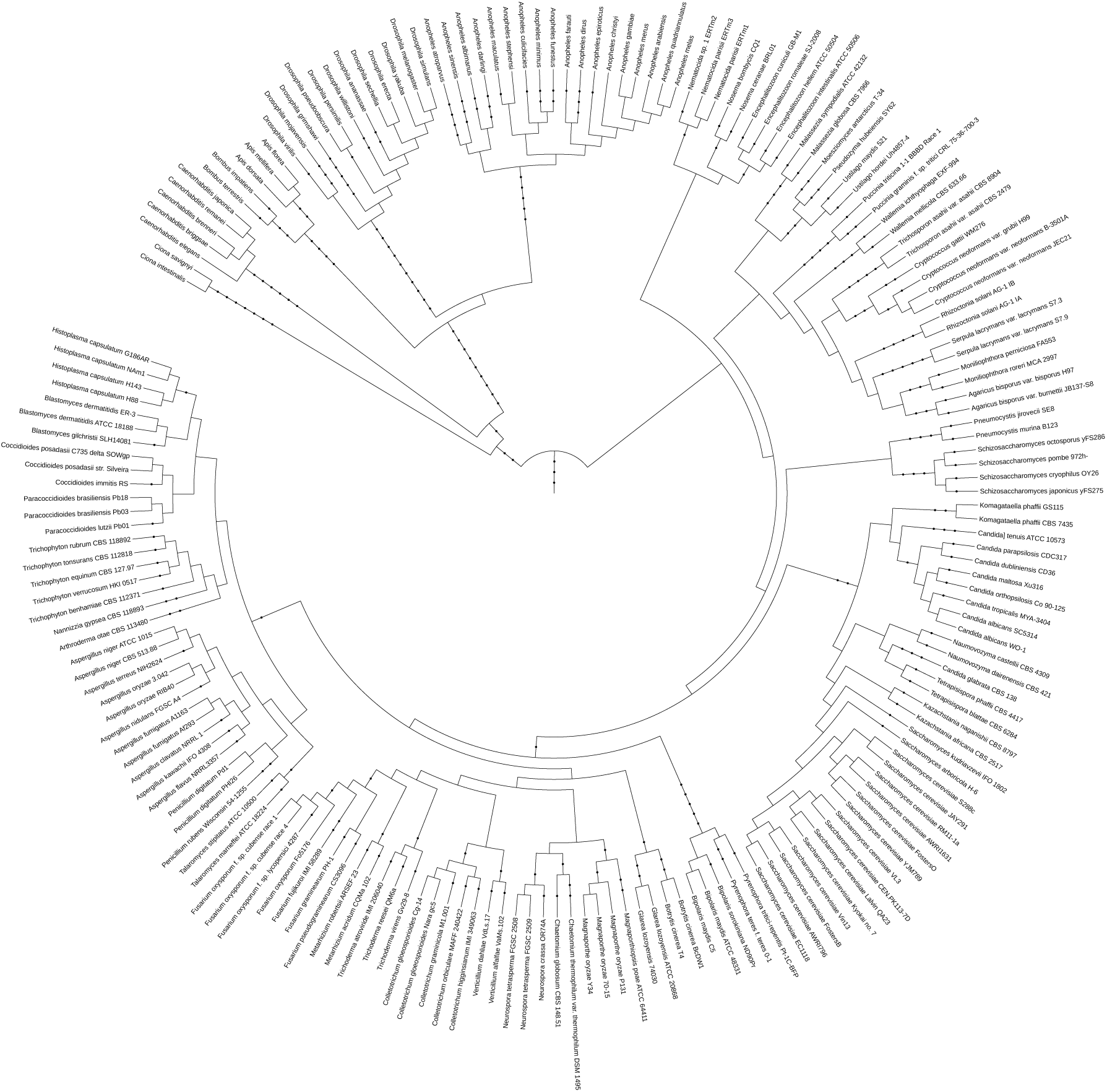
Phylogenetic tree of the _nal set of 187 eukaryotic species.

**Figure S2.**
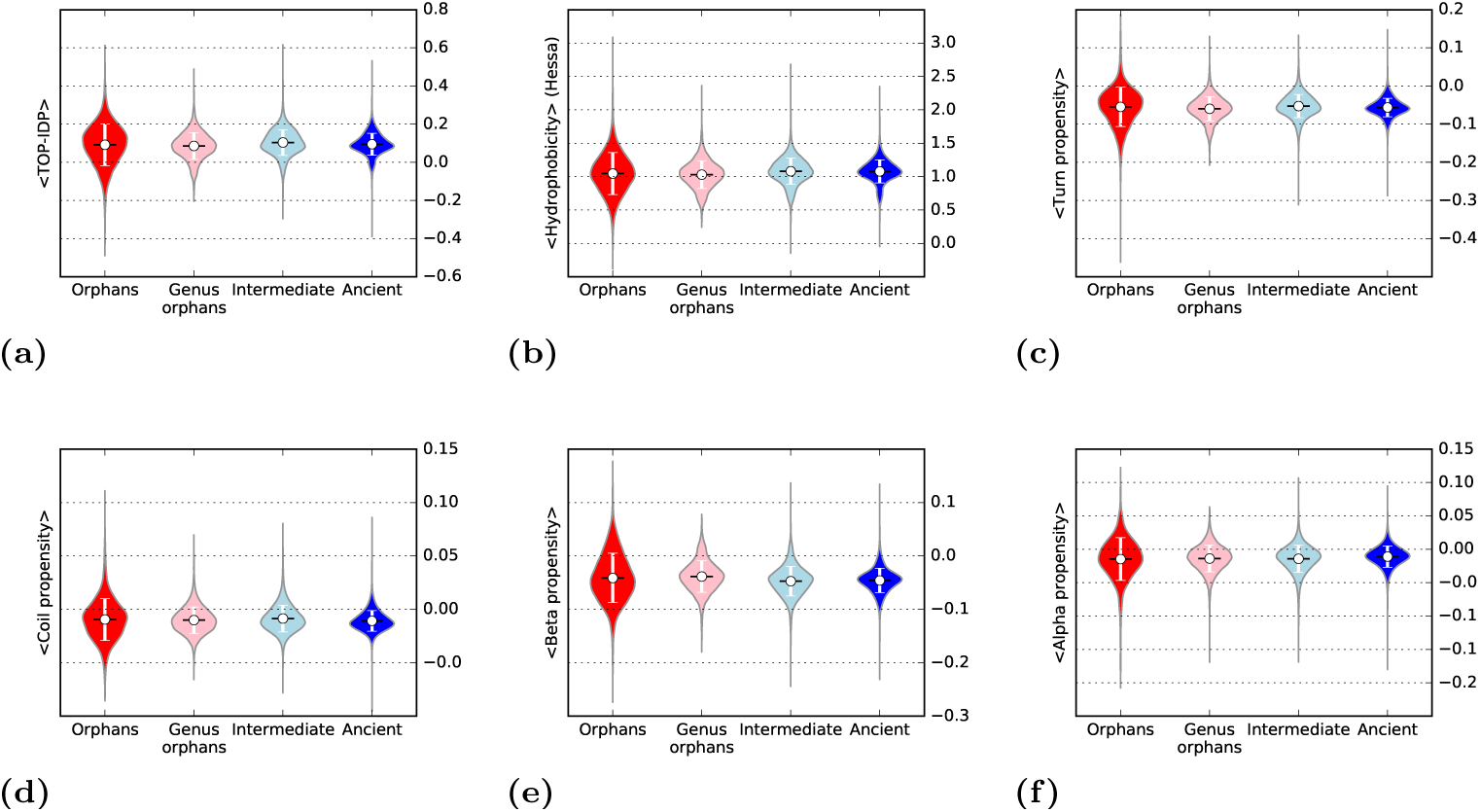
Violin plots showing several properties calculated from propensity scales, as average score. (a) Intrinsic disorder using the TOP-IDP scale, (b) hydrophobicity using the Hessa scale, (c-f) secondary structure preferences.

**Figure S3.**
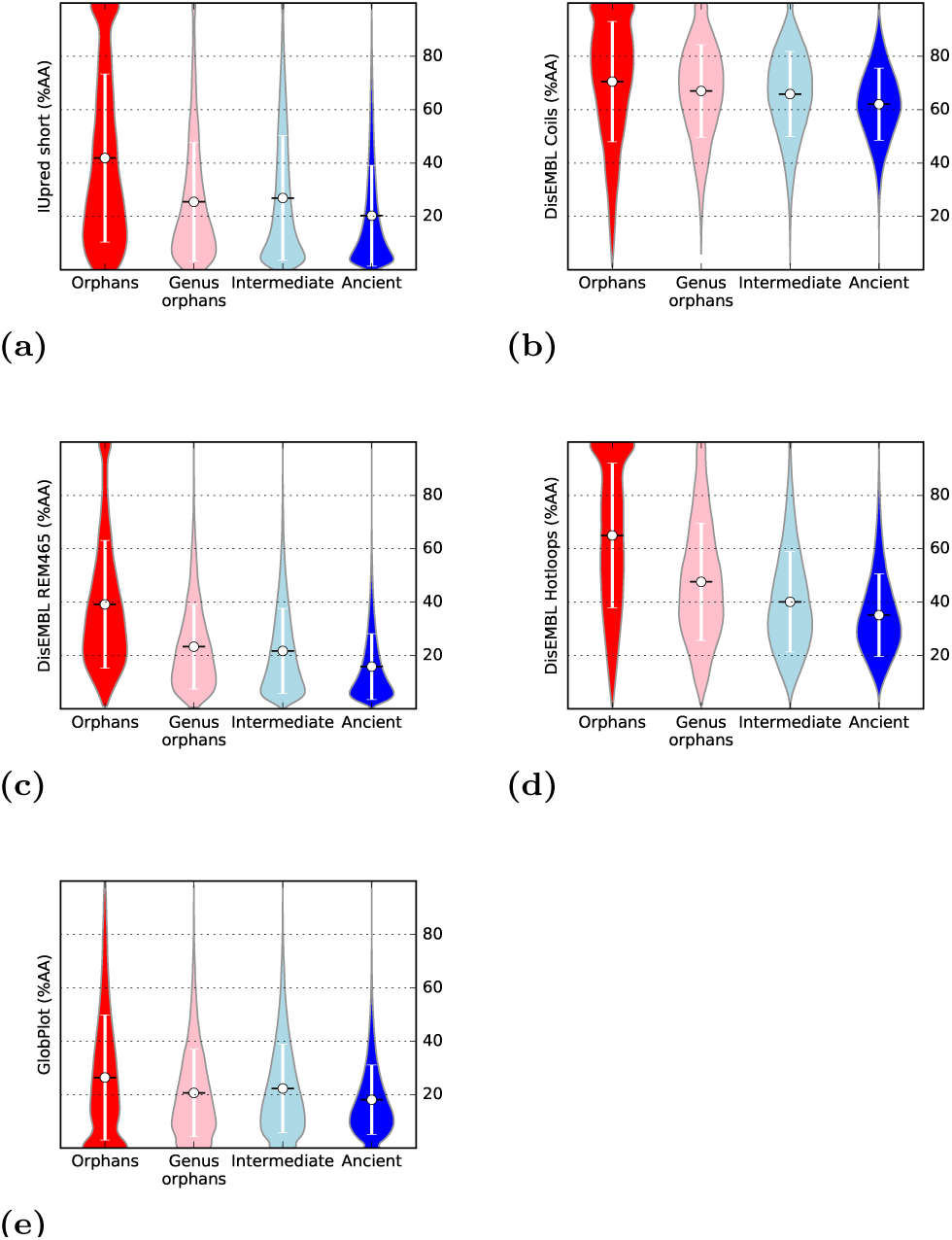
Violin plots showing intrinsic disorder as predicted by (a) IUPred (short), (b) DisEMBL Coils, (c) DisEMBL Rem465, (d) DisEMBL Hotloops and (e) GlobPlot for all the 187 species.

**Figure S4.**
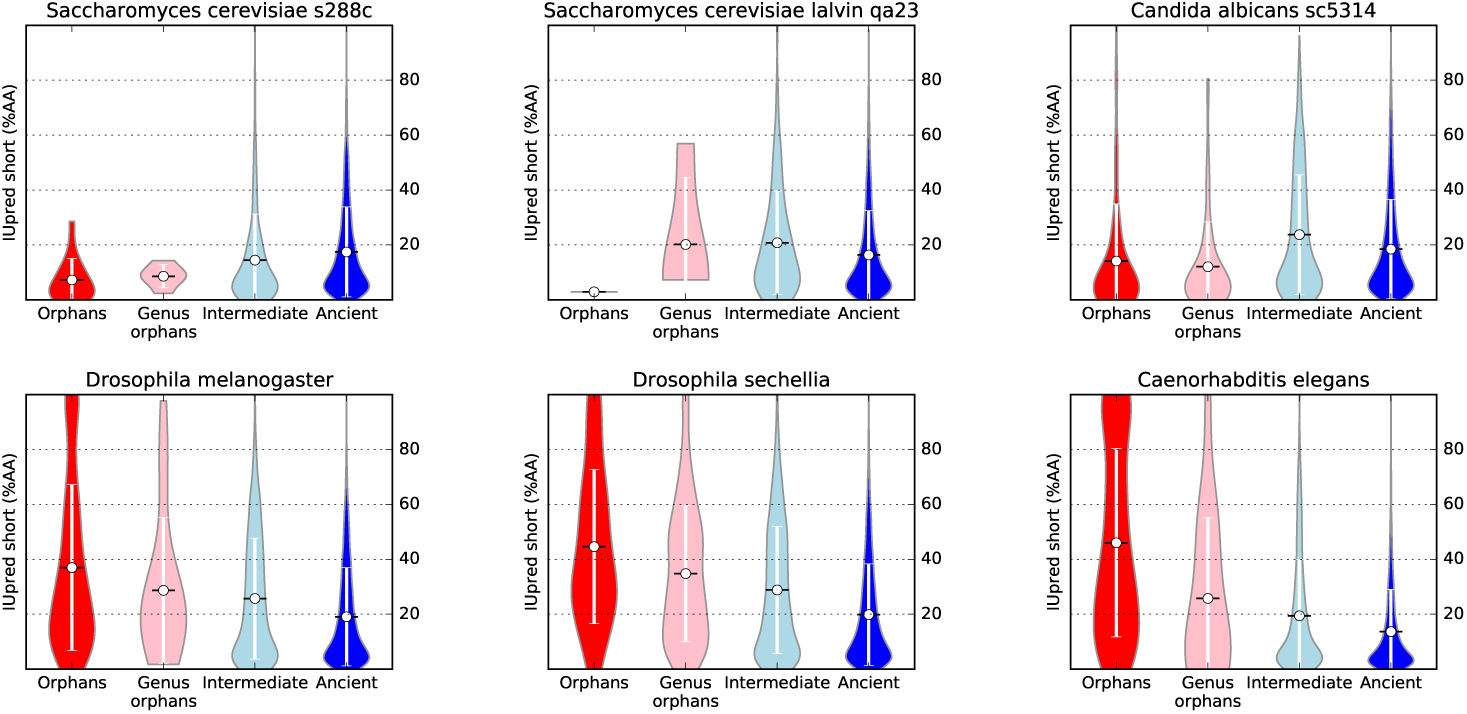
For six selected species (two strains of S. *cerevisiae*, C. *Albicans*, D. *melanogaster*, D. *sechellia* and C. *elegans*), intrinsic disorder (% of amino acid predicted as disordered by IUpred short) is shown as violin plots for proteins in the different age groups.

**Figure S5.**
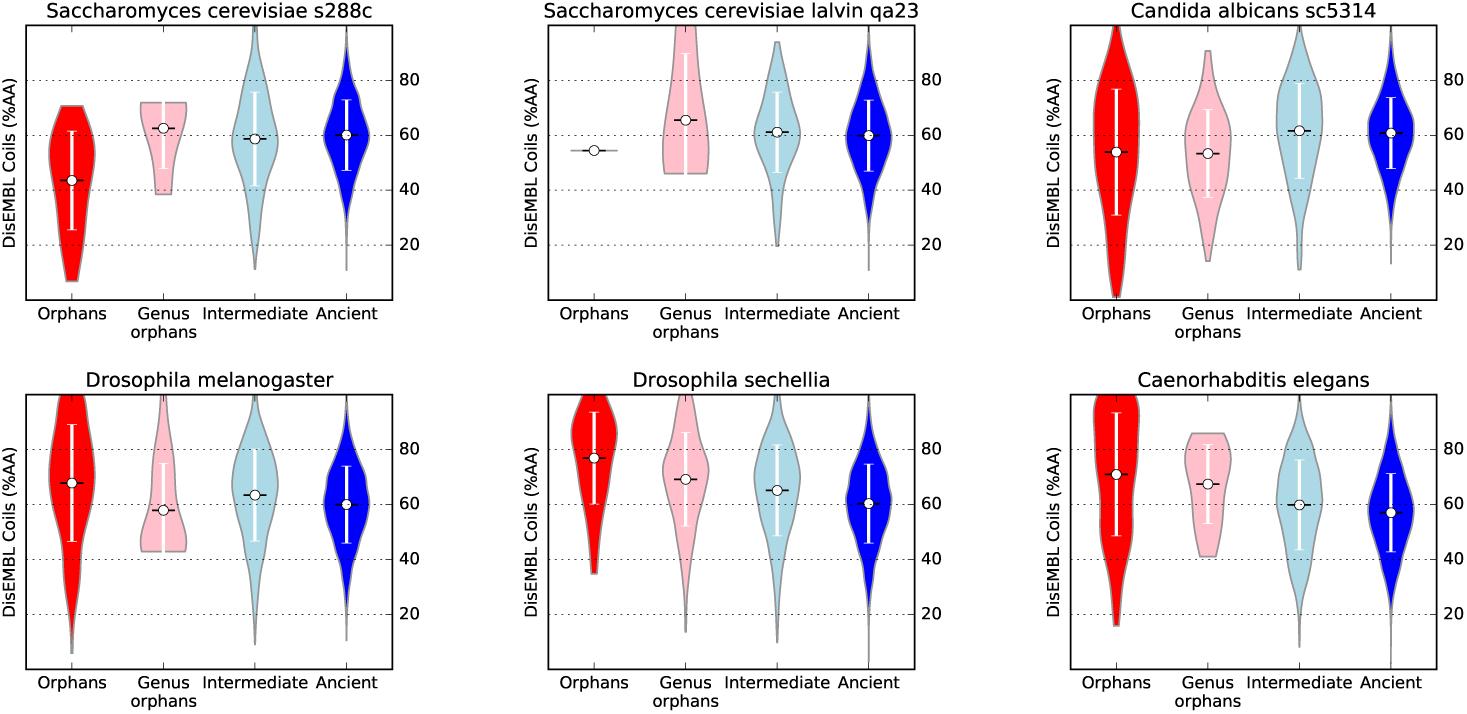
For six selected species (two strains of S. *cerevisiae*, C. *Albicans*, D. *melanogaster*, D. *sechellia* and C. *elegans*), intrinsic disorder (% of amino acid predicted as disordered by DisEMBL Coils) is shown as violin plots for proteins in the different age groups.

**Figure S6.**
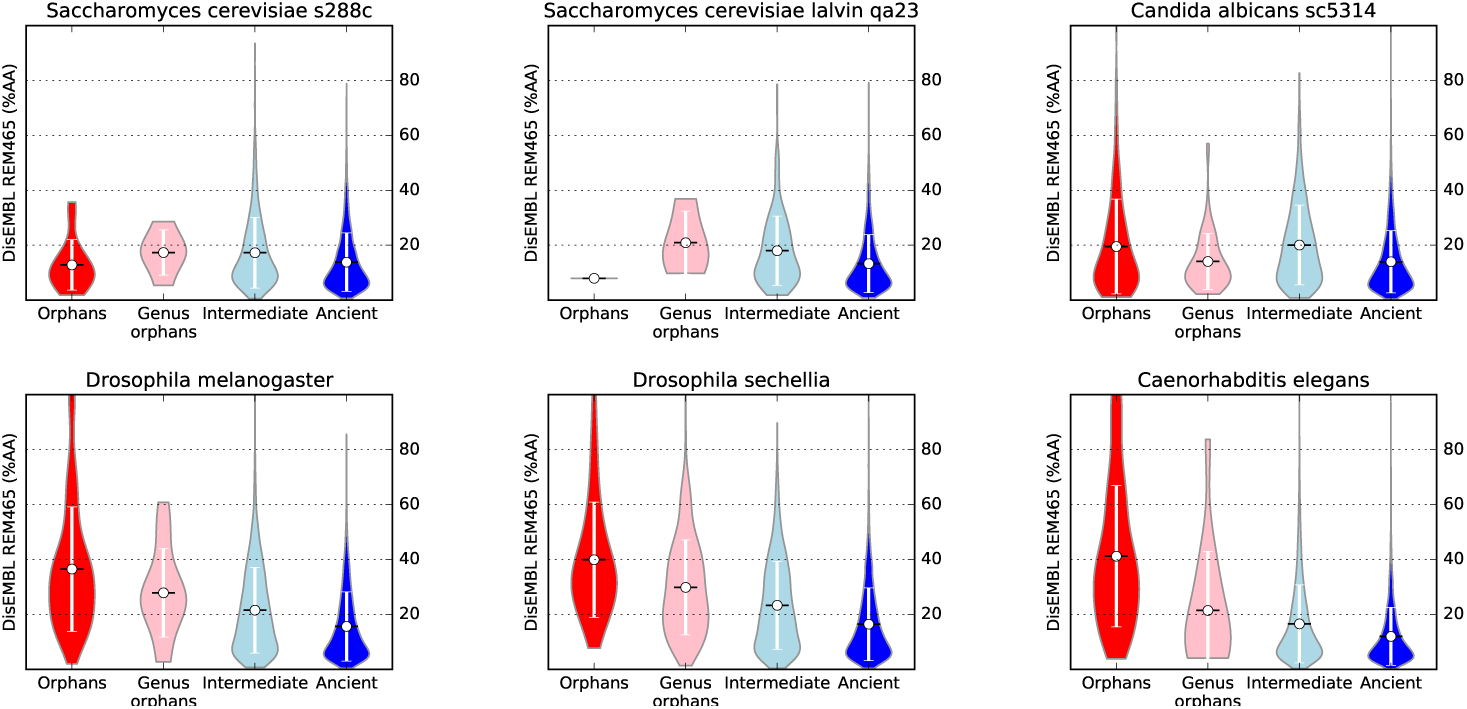
For six selected species (two strains of S. *cerevisiae*, C. *Albicans*, D. *melanogaster*, D. *sechellia* and C. *elegans*), intrinsic disorder (% of amino acid predicted as disordered by DisEMBL REM-465) is shown as violin plots for proteins in the different age groups.

**Figure S7.**
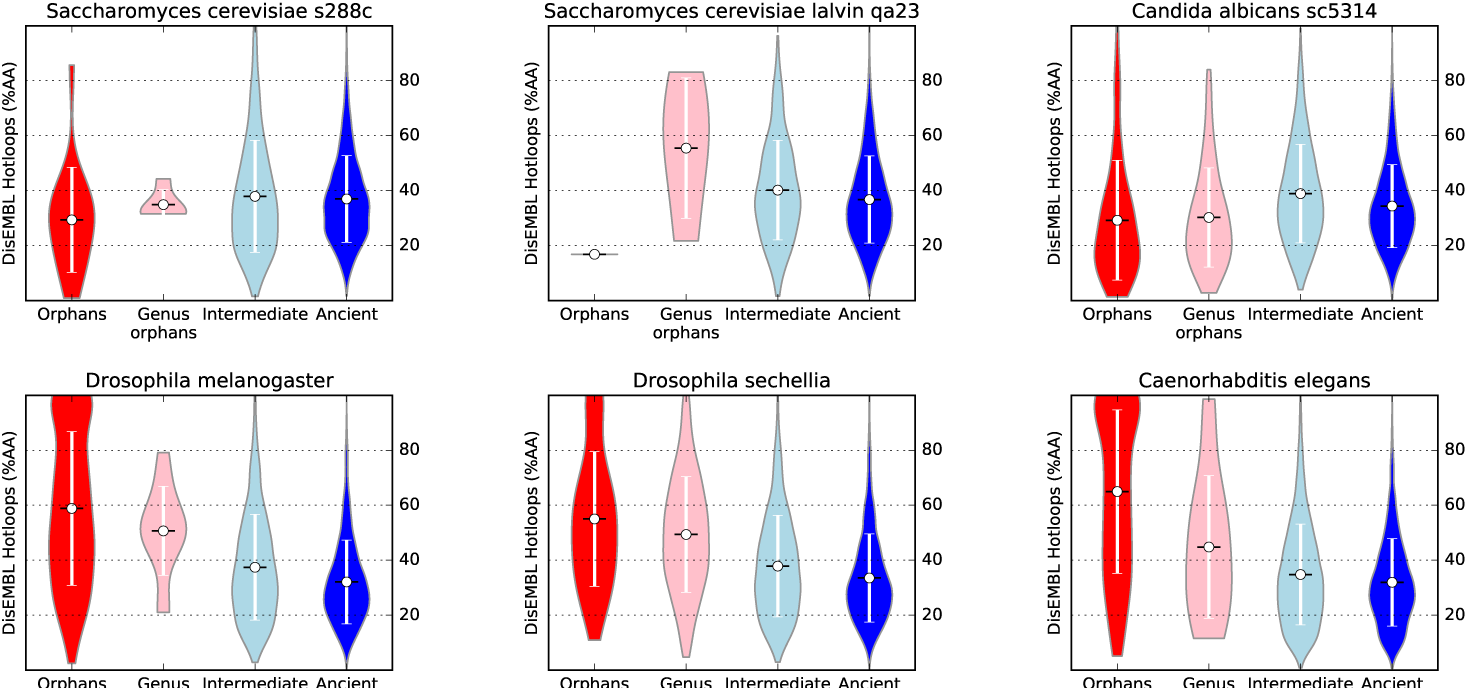
For six selected species (two strains of S. *cerevisiae*, C. *Albicans*, D. *melanogaster*, D. *sechellia* and C. *elegans*), intrinsic disorder (% of amino acid predicted as disordered by DisEMBL Hotloops) is shown as violin plots for proteins in the different age groups.

**Figure S8.**
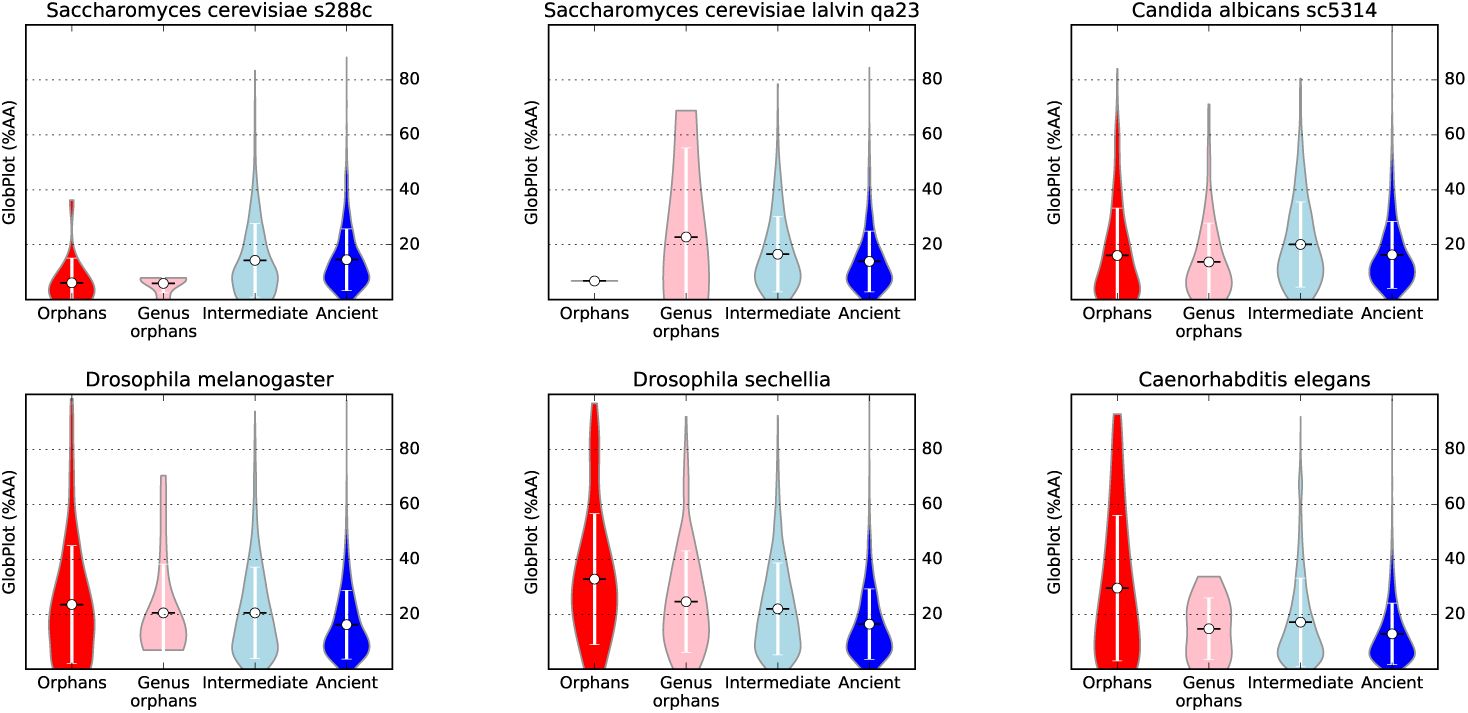
For six selected species (two strains of S. *cerevisiae*, C. *Albicans*, D. *melanogaster*, D. *sechellia* and C. *elegans*), intrinsic disorder (% of amino acid predicted as disordered by GlobPlot is shown as violin plots for proteins in the different age groups.

## References

1. Wissler L, Gadau J, Simola DF, Helmkampf M, Bornberg-Bauer E. Mechanisms and Dynamics of Orphan Gene Emergence in Insect Genomes. Genome Biology and Evolution. 2013;5(2):439–455. Available from: http://gbe.oxfordjournals.org/content/5/2/439.abstract.

2. Domazet-Loso T, Tautz D. A phylogenetically based transcriptome age index mirrors ontogenetic divergence patterns. Nature. 2010;468:815–818. Available from: http://dx.doi.org/10.1038/nature09632.

3. Tautz D, Domazet-Lošo T. The evolutionary origin of orphan genes. Nat Rev Genet. 2011 Oct;12(10):692–702. Available from: http://view.ncbi.nlm.nih.gov/pubmed/21878963.

4. Neme R, Tautz D. Evolution: dynamics of de novo gene emergence. Curr Biol. 2014 Mar;24(6):R238–40.

5. Keese PK, Gibbs A. Origins of genes: big bang or continuous creation? Proc Natl Acad Sci USA. 1992 Oct;89(20):9489–9493.

6. Siew N, Fischer D. Analysis of singleton ORFans in fully sequenced microbial genomes. Proteins. 2003 Nov;53(2):241–51. Available from: http://view.ncbi.nlm.nih.gov/pubmed/14517975.

7. Ekman D, Elofsson A. Identifying and quantifying orphan protein sequences in fungi. J Mol Biol. 2010 Feb;396(2):396–405.

8. Palmieri N, Kosiol C, Schlotterer C. The life cycle of Drosophila orphan genes. Elife. 2014;3:e01311.

9. Light S, Basile W, Elofsson A. Orphans and new gene origination, a structural and evolutionary perspective. Curr Opin Struct Biol. 2014 Jun;26:73–83.

10. Remmert M, Biegert A, Hauser A, Söding J. HHblits: lightning-fast iterative protein sequence searching by HMM-HMM alignment. Nat Methods. 2012 Feb;9(2):173–5. Available from: http://view.ncbi.nlm.nih.gov/pubmed/22198341.

11. Carvunis AR, Rolland T, Wapinski I, Calderwood MA, Yildirim MA, Simonis N, et al. Proto-genes and de novo gene birth. Nature. 2012 Jul;487(7407):370–374.

12. Neme R, Tautz D. Phylogenetic patterns of emergence of new genes support a model of frequent de novo evolution. BMC Genomics. 2013-14–117.

13. Capra JA, Williams AG, Pollard KS. ProteinHistorian: tools for the comparative analysis of eukaryote protein origin. PLoS Comput Biol. 2012;8(6):e1002567. Available from: http://view.ncbi.nlm.nih.gov/pubmed/22761559.

14. Ekman D, Bjorklund AK, Elofsson A. Quanti cation of the elevated rate of domain rearrangements in metazoa. J Mol Biol. 2007 Oct;372(5):1337–1348.

15. Wang M, Kurland CG, Caetano-Anolles G. Reductive evolution of proteomes and protein structures. Proc Natl Acad Sci U S A. 2011 Jul;108(29):11954–11958.

16. Light S, Sagit R, Sachenkova O, Ekman D, Elofsson A. Protein expansion is primarily due to indels in intrinsically disordered regions. Mol Biol Evol. 2013 Dec;30(12):2645–2653.

17. Ahrens J, Dos Santos HG, Siltberg-Liberles J. The Nuanced Interplay of Intrinsic Disorder and Other Structural Properties Driving Protein Evolution. Molecular Biology and Evolution. 2016;33(9):2248–2256. Available from: http://mbe.oxfordjournals.org/content/33/9/2248.abstract.

18. Abrusan G. Integration of new genes into cellular networks, and their structural maturation. Genetics. 2013 Dec;195(4):1407–1417.

19. Bitard-Feildel T, Heberlein M, Bornberg-Bauer E, Callebaut I. Detection of orphan domains in Drosophila using “hydrophobic cluster analysis”. Biochimie. 2015 Dec;119:244–253.

20. Kriventseva EV, Tegenfeldt F, Petty TJ, Waterhouse RM, Simão FA, Pozdnyakov IA, et al. OrthoDB v8: update of the hierarchical catalog of orthologs and the underlying free software. Nucleic Acids Res. 2015 Jan;43(Database issue):D250–6. Available from: http://view.ncbi.nlm.nih.gov/pubmed/25428351.

21. Consortium TU. UniProt: a hub for protein information. Nucleic Acids Res. 2015 Jan;43(Database issue):D204–12. Available from: http://view.ncbi.nlm.nih.gov/pubmed/25348405.

22. Letunic I, Bork P. Interactive Tree Of Life (iTOL): an online tool for phylogenetic tree display and annotation. Bioinformatics. 2007 Jan;23(1):127–128.

23. Johnson LS, Eddy SR, Portugaly E. Hidden Markov model speed heuristic and iterative HMM search procedure. BMC Bioinformatics. 2010 Aug;11:431.

24. Dosztányi Z, Csizmók V, Tompa P, Simon I. The pairwise energy content estimated from amino acid composition discriminates between folded and intrinsically unstructured proteins. J Mol Biol. 2005 Apr;347(4):827–39. Available from: http://view.ncbi.nlm.nih.gov/pubmed/15769473.

25. Linding R, Jensen LJ, Diella F, Bork P, Gibson TJ, Russell RB. Protein disorder prediction: implications for structural proteomics. Structure. 2003 Nov;11(11):1453–1459.

26. Linding R, Russell RB, Neduva V, Gibson TJ. GlobPlot: Exploring protein sequences for globularity and disorder. Nucleic Acids Res. 2003 Jul;31(13):3701–3708.

27. Bernsel A, Viklund H, Falk J, Lindahl E, von Heijne G, Elofsson A. Prediction of membrane-protein topology from first principles. Proc Natl Acad Sci U S A. 2008 May;105(20):7177–81. Available from: http://view.ncbi.nlm.nih.gov/pubmed/18477697.

28. Wootton JC, Federhen S. Analysis of compositionally biased regions in sequence databases. Methods Enzymol. 1996;266:554–71. Available from: http://view.ncbi.nlm.nih.gov/pubmed/8743706.

29. Jones DT. Protein secondary structure prediction based on position-specific scoring matrices. J Mol Biol. 1999 Sep;292(2):195–202. Available from: http://view.ncbi.nlm.nih.gov/pubmed/10493868.

30. Campen A, Williams RM, Brown CJ, Meng J, Uversky VN, Dunker AK. TOP-IDP-scale: a new amino acid scale measuring propensity for intrinsic disorder. Protein Pept Lett. 2008;15(9):956–63. Available from: http://view.ncbi.nlm.nih.gov/pubmed/18991772.

31. Hessa T, Meindl-Beinker NM, Bernsel A, Kim H, Sato Y, Lerch-Bader M, et al. Molecular code for transmembrane-helix recognition by the Sec61 translocon. Nature. 2007 Dec;450(7172):1026–1030.

32. Koehl P, Levitt M. Structure-based conformational preferences of amino acids. Proc Natl Acad Sci U S A. 1999 Oct;96(22):12524–12529.

33. Carvunis AR, Rolland T, Wapinski I, Calderwood MA, Yildirim MA, Simonis N, et al. Proto-genes and de novo gene birth. Nature. 2012 Jul;487(7407):370–4. Available from: http://view.ncbi.nlm.nih.gov/pubmed/22722833.

34. Bjorklund AK, Ekman D, Light S, Frey-Skott J, Elofsson A. Domain rearrangements in protein evolution. J Mol Biol. 2005 Nov;353(4):911–923.

35. Reeves GA, Dallman TJ, Redfern OC, Akpor A, Orengo CA. Structural diversity of domain superfamilies in the CATH database. J Mol Biol. 2006 Jul;360(3):725-741.

36. Bornberg-Bauer E, Schmitz J, Heberlein M. Emergence of de novo proteins from ‘dark genomic matter’ by ‘grow slow and moult’. Biochem Soc Trans. 2015 Oct;43(5):867–873.

37. Bernardi G. Isochores and the evolutionary genomics of vertebrates. Gene. 2000;241(1):3–17. Available from: http://www.sciencedirect.com/science/article/pii/S0378111999004850.

38. Versteeg R, van Schaik BD, van Batenburg MF, Roos M, Monajemi R, Caron H, et al. The human transcriptome map reveals extremes in gene density, intron length, GC content, and repeat pattern for domains of highly and weakly expressed genes. Genome Res. 2003 Sep;13(9):1998–2004.

39. Fickett JW. Recognition of protein coding regions in DNA sequences. Nucleic Acids Res. 1982 Sep;10(17):5303–5318.

40. Knight RD, Freeland SJ, Landweber LF. A simple model based on mutation and selection explains trends in codon and amino-acid usage and GC composition within and across genomes. Genome Biology. 2001;2(4): research0010.1. Available from: http://dx.doi.org/10.1186/gb-2001-2-4-research0010.

41. Kanaya S, Yamada Y, Kinouchi M, Kudo Y, Ikemura T. Codon usage and tRNA genes in eukaryotes: correlation of codon usage diversity with translation efficiency and with CG-dinucleotide usage as assessed by multivariate analysis. J Mol Evol. 2001 Oct-Nov;53(4–5):290–8. Available from: http://view.ncbi.nlm.nih.gov/pubmed/11675589.

